# Computational study and design of effective siRNAs to silence structural proteins associated genes of Indian SARS-CoV-2 strains

**DOI:** 10.1101/2022.02.08.479559

**Authors:** Premnath Madanagopal, Harshini Muthukumar, Kothai Thiruvengadam

## Abstract

SARS-CoV-2 is a highly transmissible and pathogenic coronavirus that first emerged in late 2019 and has since triggered a pandemic of acute respiratory disease named COVID-19 which poses a significant threat to all public health institutions in the absence of specific antiviral treatment. Since the outbreak began in March 2020, India has reported 4.77 lakh Coronavirus deaths, according to the World Health Organization (WHO). The innate RNA interference (RNAi) pathway, on the other hand, allows for the development of nucleic acid-based antiviral drugs in which complementary small interfering RNAs (siRNAs) mediate the post-transcriptional gene silencing (PTGS) of target mRNA. Therefore, in this current study, the potential of RNAi was harnessed to construct siRNA molecules that target the consensus regions of specific structural proteins associated genes of SARS-CoV-2, such as the envelope protein gene (E), membrane protein gene (M), nucleocapsid phosphoprotein gene (N), and surface glycoprotein gene (S) which are important for the viral pathogenesis. Conserved sequences of 811 SARS-CoV-2 strains from around India were collected to design 21 nucleotides long siRNA duplex based on various computational algorithms and parameters targeting E, M, N and S genes. The proposed siRNA molecules possessed sufficient nucleotide-based and other features for effective gene silencing and BLAST results revealed that siRNAs’ targets have no significant matches across the whole human genome and hence, siRNAs were found to have no off-target effects on the genome, ruling out the possibility of off-target silencing. Finally, out of 157 computationally identified siRNAs, only 4 effective siRNA molecules were selected for each target gene which is proposed to exert the best action based on GC content, free energy of folding, free energy of binding, melting temperature, heat capacity and molecular docking analysis with Human AGO2 protein. Our engineered siRNA candidates could be used as a genome-level therapeutic treatment against various sequenced SARS-CoV-2 strains in India. However, future applications will necessitate additional validations in vitro and in vivo animal models.

## 1. Introduction

COVID-19, A viral disease, caused by the new strain of severe acute respiratory syndrome coronavirus (SARS-CoV), known as SARS-CoV-2, has challenged humanity at the beginning of 2020, impacting the lives of billions of people worldwide. Since its outbreak in late December 2019 in Wuhan, China, after a sudden epidemic of atypical pneumonia with unclear illness aetiology, it has caused substantial morbidity and mortality all over the world[1]. Fever, cough, fatigue, dyspnea, and headache[2] are the most common signs of this disease, but it can also be asymptomatic[3]. Currently, molecular techniques based on the real-time reverse transcriptase-polymerase chain reaction (RT-PCR) are considered the gold standard for COVID-19 diagnosis. According to the World Health Organization (WHO), India has recorded 4.77 lakh Coronavirus deaths since the outbreak began in March 2020. Furthermore, 3.47 crore Coronavirus cases were reported in India[4]. Especially the second wave of COVID-19 has resulted in a rise in cases, a decline in crucial treatment supplies, and an increase in deaths, particularly among the young[5]. In severe cases, the patient may die due to massive alveolar damage and progressive respiratory failure[2]. Also, several occurrences of mucormycosis, popularly known as the black fungus, have been reported in patients with diabetes and patients with COVID-19, as well as patients who were recovering from infection[6].

The SARS-CoV-2 is enveloped by single-stranded positive-sense RNA and has 50% and 80% homology with Middle East Respiratory Syndrome virus and SARS-CoV, respectively and it comprises four structural proteins: envelope (E), membrane (M), nucleocapsid (N) and spike (S)[7,8]. The S, M, and E proteins are membrane-bound, while the N protein is located within the virions in complex with the genomic RNA[9]. Coronaviruses are divided into four primary genera based on their genetic makeup: Alphacoronavirus, Betacoronavirus, Gammacoronavirus, and Deltacoronavirus[10]. The first two genera primarily infect mammals, whereas the latter two primarily infect birds. Coronaviruses have genomes that are between 26 and 32 kb in size and contain 6 to 11 open reading frames (ORFs)[11]. The transmembrane trimetric spike glycoprotein extends from the viral surface and is made up of two functional subunits (S1 and S2). The S1 subunit aids in the interaction of SARS-CoV-2 with the host cell’s angiotensin-converting enzyme 2 (ACE2) receptor, whereas the S2 subunit aids in the fusion of viral and host cell membranes[12]. The M glycoprotein is the most common structural protein in coronaviruses; it spans the membrane bilayer, leaving a short NH2-terminal domain outside the virus and a long COOH terminus (cytoplasmic domain) inside[13]. All other structural proteins can bind to the M protein. Binding with M protein aids in the stabilisation of N proteins and enhances viral assembly completion by stabilising the N protein-RNA complex within the internal virion[14]. Mutations in the M protein, which cooperate with the S protein, may influence virus attachment and entrance into the host cell[15]. The E protein is the smallest and most cryptic of the major structural proteins. It is extensively expressed inside the infected cell during the replication cycle, but only a small proportion of it is incorporated into the virion envelope[16]. In the coronavirus life cycle, the RNA-binding N protein plays two important roles. Its principal function is to form the viral RNA–protein (vRNP) complex with genomic RNA and to mediate vRNP packaging into virions through poorly understood interactions between the N and M proteins. Second, the N protein is hypothesised to recruit host factors and promote RNA template switching in RTCs during the early stages of infection, facilitating viral RNA synthesis and translation[17]. Since these structural proteins (E, M, N and S) are crucial for the virus survival, these could be the main targets for designing therapeutics.

RNA interference or RNAi, the biological mechanism by which double-stranded RNA (dsRNA) induces gene silencing by targeting complementary mRNA for degradation (Figure 1)[18], is a huge breakthrough in disease therapy and is changing the way scientists analyse transcriptional activity[19] and it is a prospective tool for the control of human viral infections. Small interfering RNA (siRNA) is a double-stranded RNA of around 21 base pairs long RNA duplex bearing overhangs of two nucleotides on the 3_′_ end[20]. The strand of siRNA which has complementarity with the target gene is the guide strand and the other strand is the passenger strand. Combinations of chemically synthesised siRNA duplexes targeting SARS-CoV genomic RNA result in therapeutic activity of up to 80% inhibition, according to studies[21]. Due to the complementarity of seven nucleotides in the seed region of siRNA with the off-target gene, there is a risk of off-target gene silence or unexpected gene downregulation. The off-target binding impact is mediated by the melting temperature or thermodynamic stability of the duplex of seed siRNA sequence (2-8 nucleotides of siRNA guide strand from 5′ end) and the target gene, according to studies. Thus, a siRNA with perfect complementarity solely for the target gene and low seed-target duplex thermostability (Tm less than 21.5°C) can efficiently remove siRNA off-target binding. Furthermore, selecting a siRNA with at least two mismatches with any other off-target region can lower the likelihood of siRNA binding to the unwanted off-target sequence[22,23]. Along with the seed region, the non-seed region of the guide strand has been shown to be effective in mediating off-target impact, yet there was a negative correlation between Tm value and GC content and off-target effect[24].

**Figure 1:**
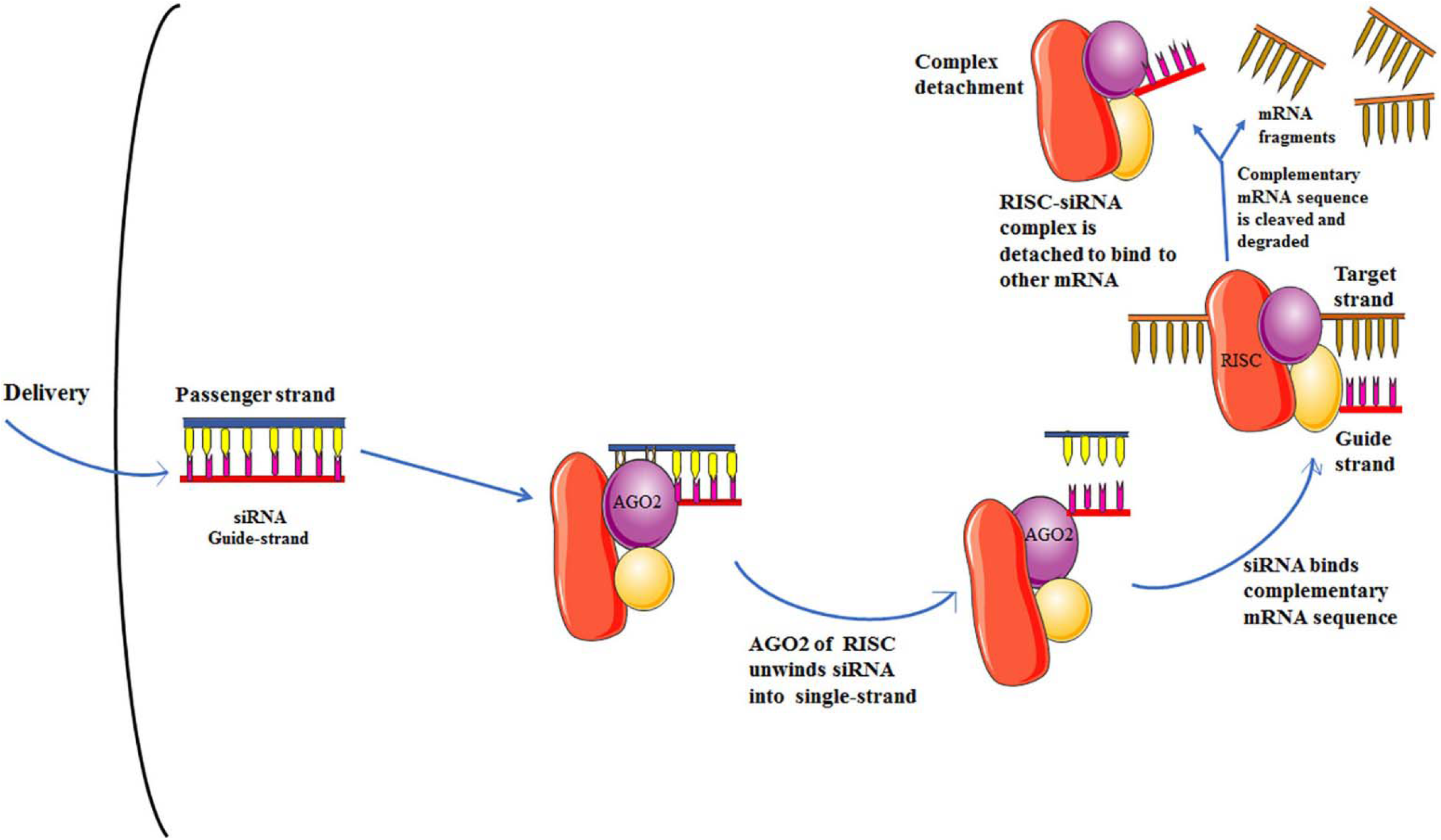
Graphical representation of the siRNA-mediated gene silencing mechanism.

Because structural proteins expressed by the E, M, N and S genes are involved in virus survival and infectivity, they can be used as a target for suppressing SARS-CoV-2 infection. *In silico* approaches to discovering innovative therapeutic approaches are becoming increasingly popular, and viruses are no exception. Therefore, in this present study, we have designed siRNAs specific to the various conserved region of the envelope protein, membrane glycoprotein, nucleocapsid and surface glycoprotein genes by analyzing 811 Indian SARS-CoV-2 strains and finally shortlisted 4 effective siRNA molecules for each gene which will inhibit the translations of these target genes and allow the host to discard this infection (Figure 2).

**Figure 2.**
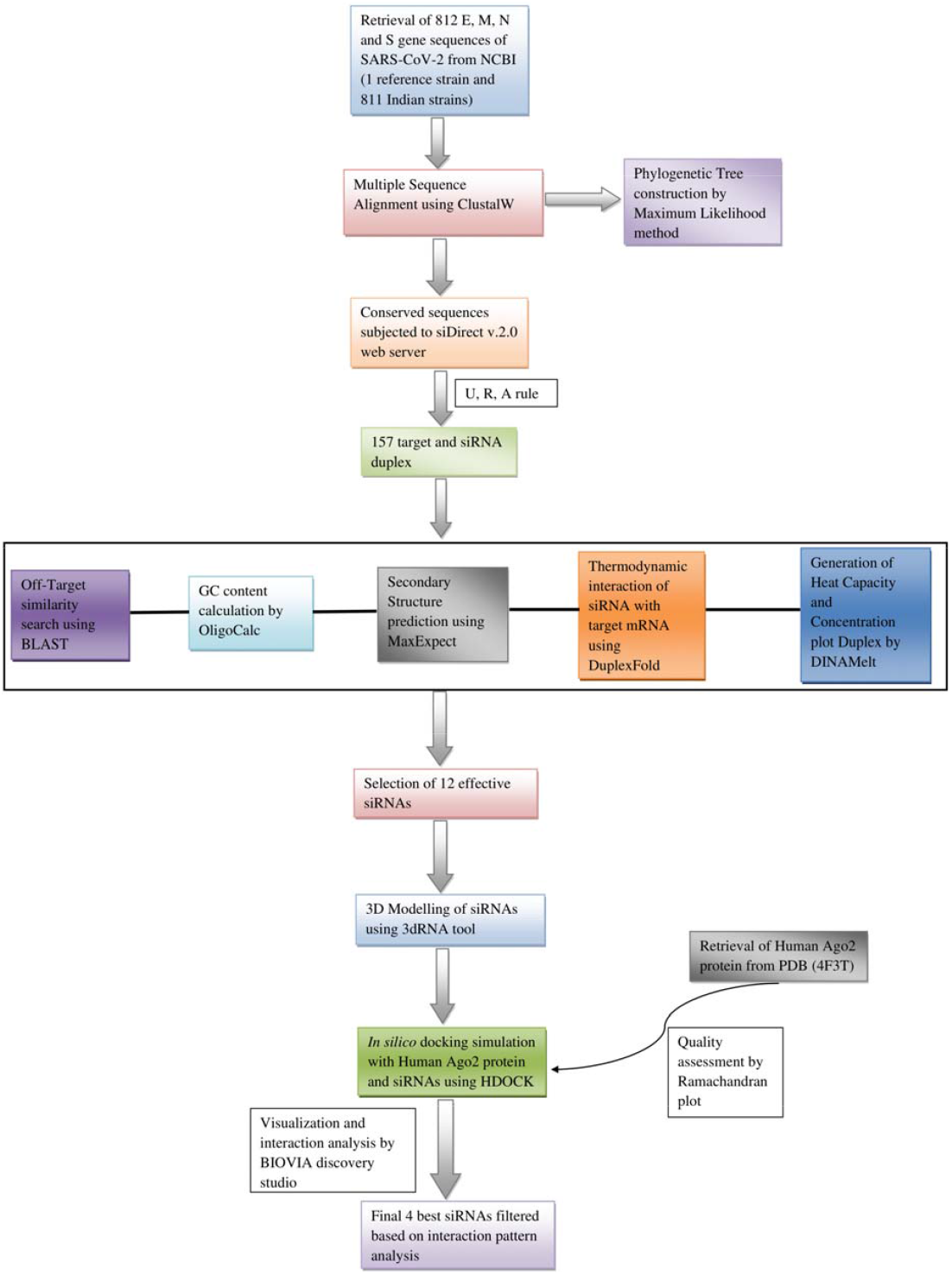
Flowchart depicting the workflow of the methodology used in the study.

## 2. Materials and Methods

### 2.1. Retrieval of gene sequences

812 gene sequences of the envelope gene(E), membrane gene(M), nucleocapsid phosphoprotein gene(N) and surface glycoprotein gene(S) of various Indian SARS-CoV-2 strains were retrieved from the GenBank available at the National Center for Biotechnology Information (NCBI) (Supplementary Table 1)[25]. Out of 812 gene sequences, 811 were Indian strains and the remaining one was the Wuhan-Hu-1 strain (China, Genbank accession: NC_045512.2) that was used as a reference genome sequence for multiple sequence alignment and site numbering for amino acids.

### 2.2. Multiple sequence alignment

Multiple sequence alignment of all 812 gene sequences of E, M, N and S genes were performed using the ClustalW[26] algorithm in the MEGA-X[27] program to find the conserved regions.

### 2.3. Target recognition and potential siRNA designing

siDirect version 2.0[23] is used to design effective and target-specific siRNA molecules against SARS-CoV-2 E, M, N and S gene sequences. This tool utilizes some rules such as Ui-Tei[28], Amarzguioui[29] and Reynolds[30] algorithms (Table 1) for designing siRNAs and the melting temperature (Tm) of the seed-target duplex was kept below 21.5°C as a default parameter. To avoid off-target silencing, it chooses siRNA sequences with at least two mismatches to any other non-targeted transcripts.

**Table 1.** Rules/Algorithms for designing of effective siRNA molecules.

| Ui-Tei rules | Amarzguioui rules | Reynolds rules |
| --- | --- | --- |
| • A/U at the 5' end of the antisense strand<br>• G/C at the 5' end of the sense strand<br>• At least five A/U residues in the 5' terminal one-third of the antisense strand<br>• The absence of any GC stretches of more than 9 nt in length | • The A/U differential of the duplex end should be >0<br>• Strong binding of 5 sense strand<br>• Position 1 must have any bases other than U<br>• Position 6 must have A constantly<br>• Weak attachment of 3' sense/passenger strand. | • The designed siRNA must maintain a GC content between 30% to 52% (1 point)<br>• Occurrence of three or more A/U base pair at position 15–19 of sense strand (Each A/U base pair in this region earns one point)<br>• The T<sub>m</sub> (melting temperature) of designed siRNA must be greater than –20 °C (1 point)<br>• Position 19 of sense strand must contain A (1 point)<br>• Position 3 of sense strand must include A (1 point)<br>• Position 10 of the sense strand should have U (1 point)<br>• Position 13 of the sense strand must contain any bases other than G (1 point)<br>• Threshold for efficient siRNAs score = 6 |

### 2.4. Off-target investigation using BLAST

The BLAST[31] tool was used to identify the possible off-target matches in a human genomic transcript against the whole GenBank[32] database by applying the expected threshold value 10 and BLOSUM 62 matrix[33] as a parameter.

### 2.5. GC content calculation

OligoCalc[34] was used to analyze the GC content of predicted siRNA molecules.

### 2.6. Secondary structure prediction

The secondary structure and the free energy (ΔG) of folding for predicted siRNA molecules were computed using the MaxExpect[35] program of the RNA structure web server[36]. Higher energy values imply that those molecules are less likely to fold and it indicates that they are better candidates.

### 2.7. Calculation of RNA–RNA Interaction Through Thermodynamics

The thermodynamic interaction between the target strand and the siRNA guide strand was predicted using the DuplexFold[37] program of the RNA structure web server[36]. It folds two sequences of RNA into their lowest hybrid free energy conformation. Higher interaction between the target and the guide strand will aid in a better predictor of siRNA effectiveness.

### 2.8. Calculation of heat capacity and concentration plot duplex

The heat capacity plot and concentration plot were calculated for the predicted siRNAs using the DINA Melt web server[38]. The ensemble heat capacity (Cp) is plotted as a function of temperature, with the melting temperature Tm (Cp) indicated in Table 2. The detailed heat capacity plot also shows the contributions of each species to the ensemble heat capacity, whereas the concentration plot- Tm (Conc), the point at which the concentration of double-stranded molecules of one-half of its maximum value defines the melting temperature Tm (Conc).

**Table 2.**
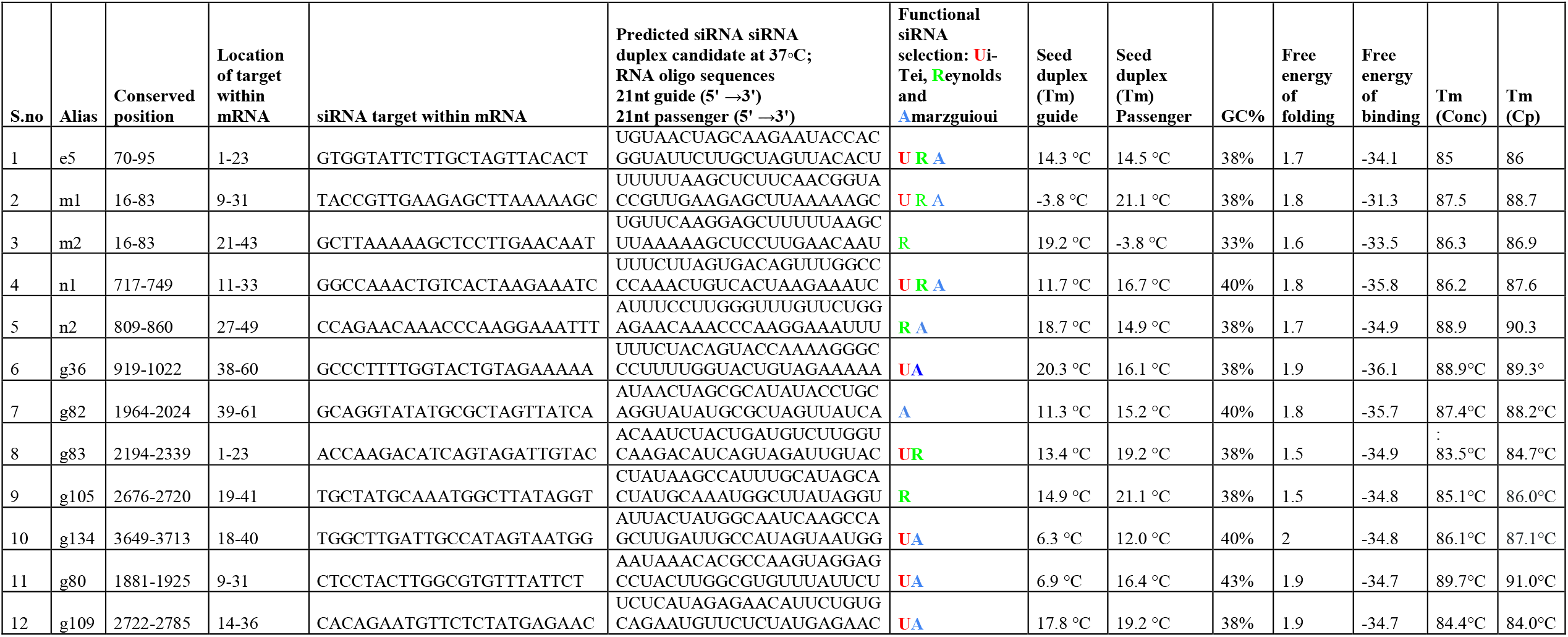
Effective siRNA molecules with GC%, free energy of folding and free energy of binding with target.

### 2.9. RNA modelling and Protein preparation

The proper interaction between siRNA duplex and RISC complex, as well as guide siRNA strand and target mRNA inside RISC complex, are required to trigger a sufficient antiviral response via RNAi-mediated viral gene silencing[18]. Hence for interaction pattern analysis, molecular docking was carried over between siRNA (guide strand) and argonaute protein. The 3D modelled structure of guide siRNA was generated using a 3dRNA v2.0 web server[39]. 3dRNA is an automatic, fast, and high-accuracy RNA tertiary structure prediction method. It uses sequence and secondary structure information to build three-dimensional structures of RNA from template segments. We used an optimization procedure which is an integrated 3dRNA pipeline, to model our predicted siRNAs.

The three-dimensional structure of human argonaute 2 (Ago2) protein (Figure 3a) was retrieved from RCSB Protein Data Bank (PDB ID: 4F3T)[40]. BIOVIA discovery studio[41] was used to prepare the proteins through correction of bonds, removal of unrelated chemical complexes, elimination of water molecules and hetatom groups. The missing side-chain atoms in protein have been reconstructed and then the structure was minimized with the partial implementation of the GROMOS96 force-Field using Swiss-PdbViewer[42]. The quality of the model was checked using Ramachandran Plot generated by PROCHECK[43].

**Figure 3.**
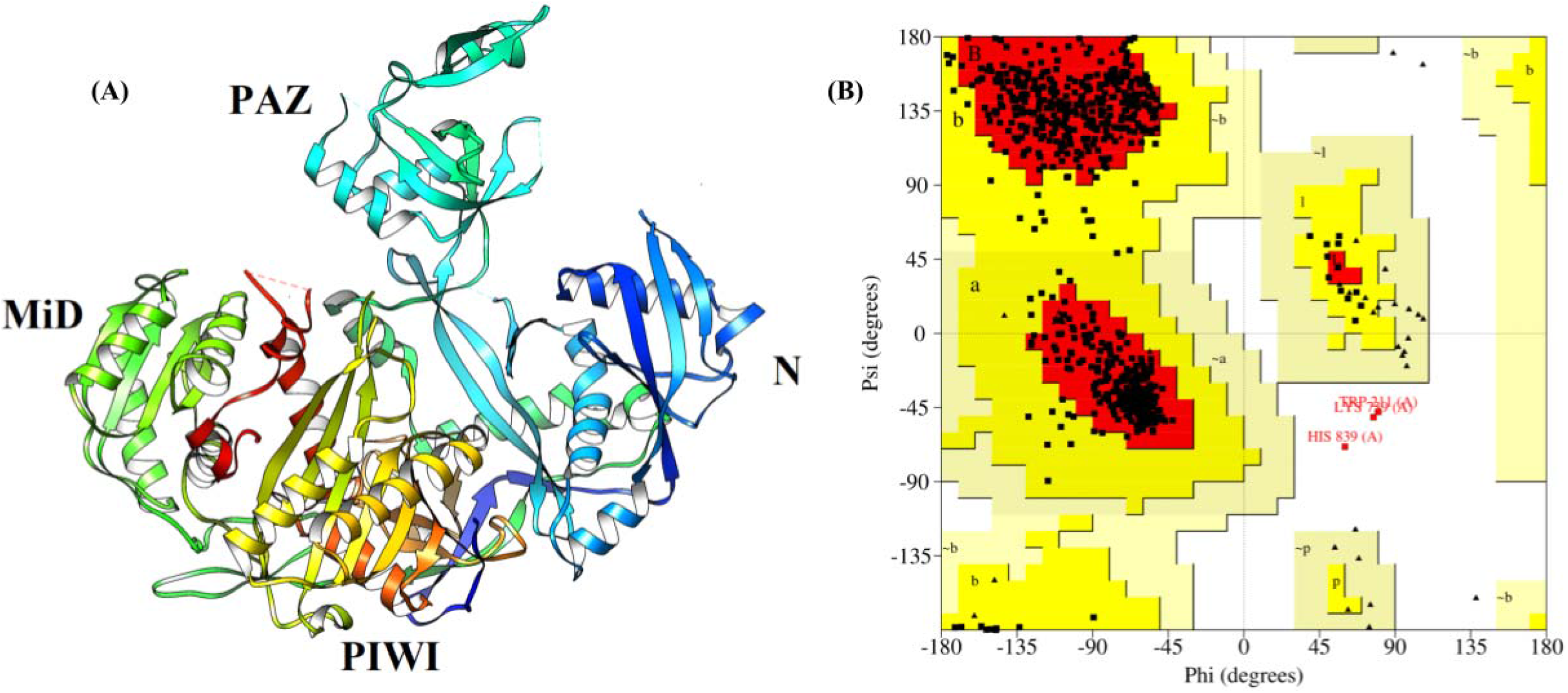
A) The cartoon representation of the structure of Human Argonaute 2 (Ago2) protein. B) The Ramachandran plot of Human Ago2 protein.

### 2.10. Molecular Docking and Interaction analysis

The molecular docking between siRNA (guide strand) and argonaute protein was performed using the HDOCK server[44]. This server uses a hierarchical Fast Fourier Transform (FFT) - related docking program. It is a hybrid docking algorithm that uses free docking and template-based modelling and shows high efficacy and robustness. Further docking analysis of protein-siRNA complex was performed using BIOVIA discovery studio[41].

## 3. Results

### 3.1. Target specific designing of potential siRNA molecules

siDirect 2.0 web server predicted 7 siRNA for the envelope, 10 siRNA for membrane, 3 siRNA for nucleocapsid phosphoprotein and 137 siRNA for surface glycoprotein (Supplementary Table 6, 7, 8 and 9) that followed algorithms of Ui-Tei, Amarzguioui and Reynolds (Table 1). The seed target duplex stability (Tm) values for all predicted siRNAs were less than 21.5 °C, which shows that predicted siRNAs may avoid nontarget binding.

### 3.2. Analysis of target and off-target exclusion using blast

The consensus targets obtained by siDirect 2.0 were subjected to NCBI-BLAST for searching similarity against the whole human genome, however, no significant matches were found. This shows that our predicted siRNA may not interact in any place other than the viral target site.

### 3.3. GC content calculation of predicted siRNA

The sequence’s GC content is a vital parameter that influences siRNA functionality. It has been suggested to select sequences with a low to medium proportion (31.6 to 57.9%) of GC content[45]. GC content analysis of the predicted siRNAs was ranged from 21% to 38% for the envelope, 10% to 38% for membrane, 36% to 40% for nucleocapsid phosphoprotein and 10% to 43% for surface glycoprotein (Supplementary Tables 10, 11, 12 and 13).

### 3.4. Secondary structure determination

The probable folding and minimum free energy of folding of the predicted siRNA guide strands were calculated with the assumption that siRNA with the positive free energy of folding may be more permissible to the target, resulting in better gene silencing[46]. The calculated free energy of folding ranged from 1.3 to 1.7 for the envelope, from 1.6 to 2 for membrane, from 1.7 to 1.9 for nucleocapsid phosphoprotein and from 1.1 to 2.9 for surface glycoprotein (Supplementary Tables 10, 11, 12 and 13).

### 3.5. Thermodynamics of target-guide strand interaction

The binding free energy between the target and guide strand was calculated. The values spanned from -25.8 to -34.1 for envelope, -20.9 to -33.5 for membrane, -32.5 to -35.8 for nucleocapsid phosphoprotein and -20.5 to -36.1 for surface glycoprotein (Supplementary Tables 10, 11, 12 and 13).

### 3.6. Determination of heat capacity and duplex concentration plot

The predicted siRNAs’ Tm(Cp) and Tm(Conc) were computed. The greater the values of these two melting temperatures, the more effective the siRNA species. Tm(Conc) values ranged from 76.4 °C to 85 °C for envelope, 66.8 °C to 87.5 °C for membrane, 86.2 °C to 88.9 °C for nucleocapsid phosphoprotein and 63.4 °C to 91.1 °C for surface glycoprotein. Tm(Cp) values ranged from 77.6 °C to 86 °C for envelope, 68.4 °C to 88.7 °C for membrane, 87.6 °C to 89.4 °C for nucleocapsid phosphoprotein and from 69.0 °C to 92.1 °C for surface glycoprotein (Supplementary Tables 10, 11, 12 and 13).

### 3.7. Analysis of molecular modelling and docking of best siRNA (guide strand) & Ago2

According to the Ramachandran plot of Ago2 protein, the percentage of residues that resided in the core region, allowed region, moderately allowed region, and disallowed region was 90.7%, 8.9%, 0.0%, and 0.4% respectively (Figure 3b). Best siRNAs have been filtered for each gene based on the criteria as follows GC content (33% to 43%) and Free energy of binding (>=-31.3) (Table 2). The docking score, interaction statistics and interactive residues were provided in Table 3. The docking score ranged from -290.15 to -358.89 where g82 had the lowest docking score and g80 had the highest binding affinity.

**Table 3.** siRNAs docking score and interaction statistics.

| S.no | siRNA in Ago2-siRNA complex | HDock Docking score | Interaction Category |  |  |  | Total number of interacting residues |
| --- | --- | --- | --- | --- | --- | --- | --- |
|  |  |  | Salt bridge (Hydrogen Bond+Electrostatic) | Hydrogen bonds | Electrostatic bonds | Hydrophobic bonds |  |
| 1 | e5 | -322.34 | 7 | 16 | 8 | 5 | 23 |
| 2 | m1 | -324.56 | 3 | 13 | 11 | 6 | 27 |
| 3 | m2 | -348.34 | 3 | 15 | 8 | 3 | 20 |
| 4 | n1 | -335.64 | 1 | 15 | 7 | 0 | 16 |
| 5 | n2 | -294.72 | 1 | 10 | 9 | 5 | 14 |
| 6 | g36 | -354.19 | 1 | 12 | 7 | 2 | 16 |
| 7 | g82 | -290.15 | 1 | 21 | 10 | 3 | 19 |
| 8 | g83 | -326.1 | 1 | 12 | 4 | 5 | 12 |
| 9 | g105 | -312.45 | 2 | 14 | 3 | 3 | 17 |
| 10 | g134 | -338.34 | 0 | 14 | 3 | 2 | 12 |
| 11 | g80 | -358.89 | 4 | 8 | 10 | 2 | 17 |
| 12 | g109 | -319.69 | 3 | 12 | 7 | 2 | 12 |

## 4. Discussion

COVID-19, respiratory illness caused by the SARS-CoV-2 coronavirus, has recently become pandemic. The pandemic is sweeping through India at a pace that has staggered scientists. To combat this pandemic, promising varied techniques such as gene therapy, as well as other therapeutic tactics such as the creation of vaccines, drugs, monoclonal antibodies, peptides, and other therapeutic strategies are being suggested. The present study introduced an *in silico* approach to design potential siRNA molecules of Indian SARS-CoV-2 strains targeting virus structural genes which included envelope protein (E), membrane protein (M), nucleocapsid phosphoprotein (N) and surface glycoprotein (S).

Among the 811 SARS-CoV-2 strains from throughout India, 243 conserved regions (10 envelope protein, 26 membrane protein, 67 nucleocapsid phosphoprotein and 140 surface glycoprotein) (Supplementary Tables 2, 3, 4 and 5) were found. Conserved regions that are less than 21 nucleotides were taken out of the further analysis. The siDirect webserver was used to identify potential targets and generate corresponding siRNAs using conserved sequences. The task is completed by siDirect in three steps: highly functional siRNA selection, seed-dependent off-target effects reduction, and near-perfect matched genes removal. There were 157 target regions, 7 envelope proteins, 10 membrane proteins, 3 nucleocapsid phosphoproteins, and 137 surface glycoproteins (Supplementary Tables 6, 7, 8 and 9). To improve the results, the U, R, A (Ui-Tei, Amarzguioui, and Reynolds) guidelines (Table 1) were used to identify the siRNAs. The formation of non-target siRNA bonds was eliminated by lowering the melting temperature (Tm) below 21.5 degrees Celsius.

Despite the fact that the primary goal of siRNA design is to silence specific targets, there is still the possibility of silencing an unknown number of unwanted genes[47]. The off-target activity caused by siRNA can be explained by two mechanisms: mRNA depletion or translational suppression at the protein level[48]. First, siRNA can tolerate several mismatches to the mRNA target while also retaining good silencing capacity with imperfect complementarity[47]. Another possible mechanism for off-target activity is that siRNAs’ physical structure, ∼21 nucleotides (nt) RNA oligomers, appears to be identical to the related class of microRNAs. MicroRNAs are short, endogenously transcribed RNAs that prevent mRNA from being translated rather than being degraded. MicroRNAs appear to contain RNA oligo and RNA target mismatches built into their structure. Together the mechanisms of siRNA mismatch tolerance and microRNA translation inhibition create a risk of off-target activity when ∼21 nt RNAs are introduced into human cells[47,48]. As a result, off-target effects may occur even when siRNAs are generated using different algorithms. To ensure siRNA target specificity, siRNA design processes are often followed by a BLAST search for cross-reactive 21-bp siRNA sequences. BLAST results show that the SARS-CoV-2 target sequences (E, M, N, and S) had no significant hits against the entire genome, ruling out the possibility of off-target silencing.

The GC-content of siRNA influences its functioning, and there is an inverse link between GC-content and siRNA function. siRNA targets with a very high GC content may be more prone to folding, which could weaken target accessibility. For a siRNA to be effective, it needs to have a medium proportion of GC content, which ranges from 31.6 to 57.9%[45]. Therefore, molecules with a GC content of less than ∼32% were excluded from the final selection. For each of the 157 predicted species, the GC content ranged from 10% to 43%. The GC content of the filtered siRNAs is greater than or equivalent to 33%. (Table 2).

The RNA structure webserver was used to estimate probable folding structures and corresponding minimal free energy using guide strands of predicted siRNAs. At 37°C, the free energy of folding of the final siRNAs is more than zero (Figure 4, Table 2), implying that the predicted siRNAs are more accessible for efficient binding.

**Figure 4.**
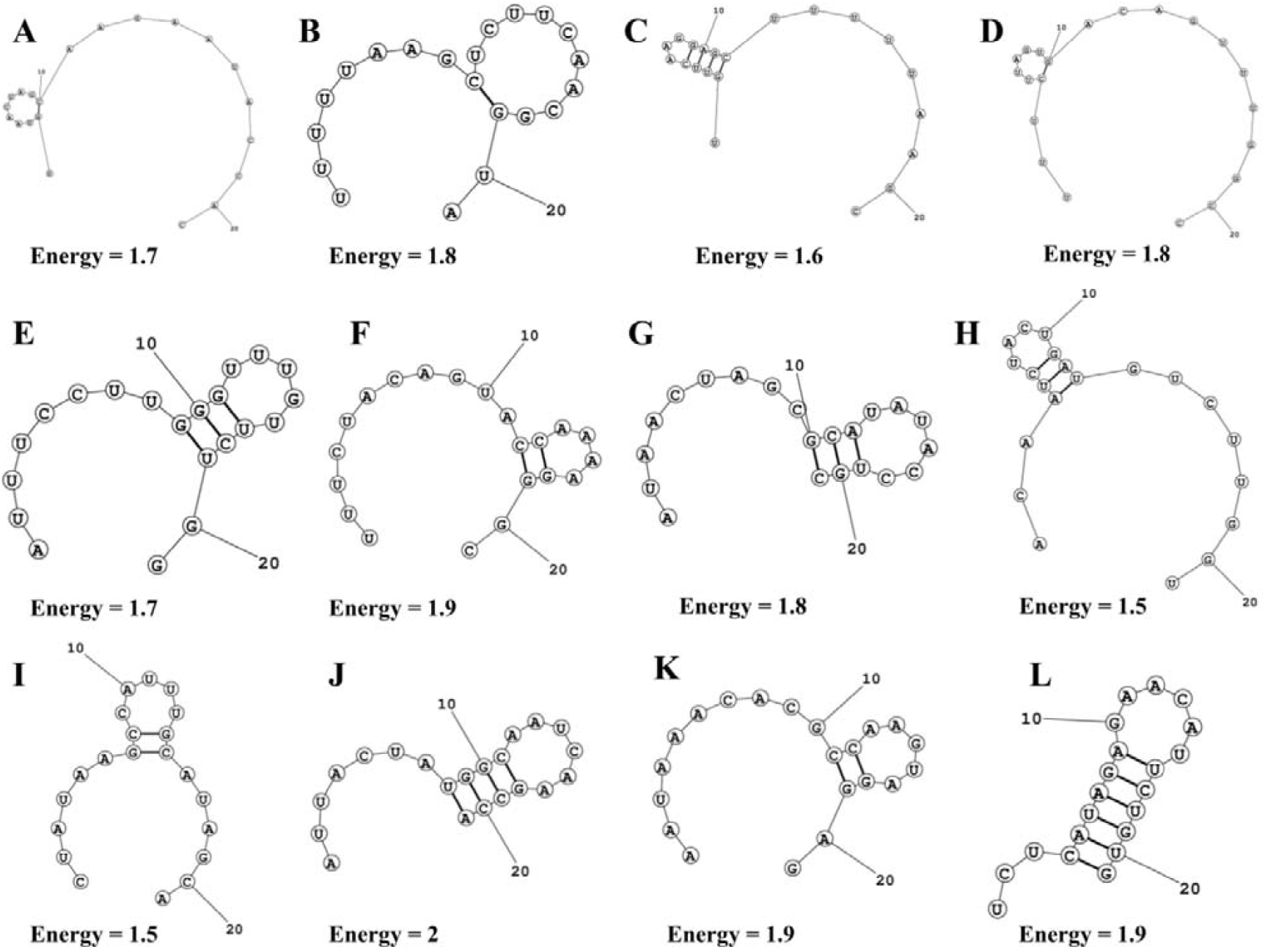
The possible folding and minimum free energy of the guide strands of the predicted siRNA molecules. The structures are for A. e5 B. m1 C. m2 D. n1 E. n2 F. g36 G. g82 H. g83 I. g105 J. g134 K. g80 L. g109 siRNAs.

DuplexFold was used to calculate the binding energy of the target mRNA and guide siRNA interactions. Lower binding energy suggests a better interaction, and hence a better likelihood of inhibiting the target. All 157 predicted siRNAs had binding free energy values ranging from -36.1 to -20.5. (Supplementary Tables 10, 11, 12 and 13). We chose the best siRNAs for each E, M, S, and N gene based on GC content and free energy of binding criteria for further docking analysis. The selected siRNAs have a free energy of binding equal to or less than - 31.3 (Figure 5, Table 2), implying that predicted siRNAs are more interactive with their targets.

**Figure 5.**
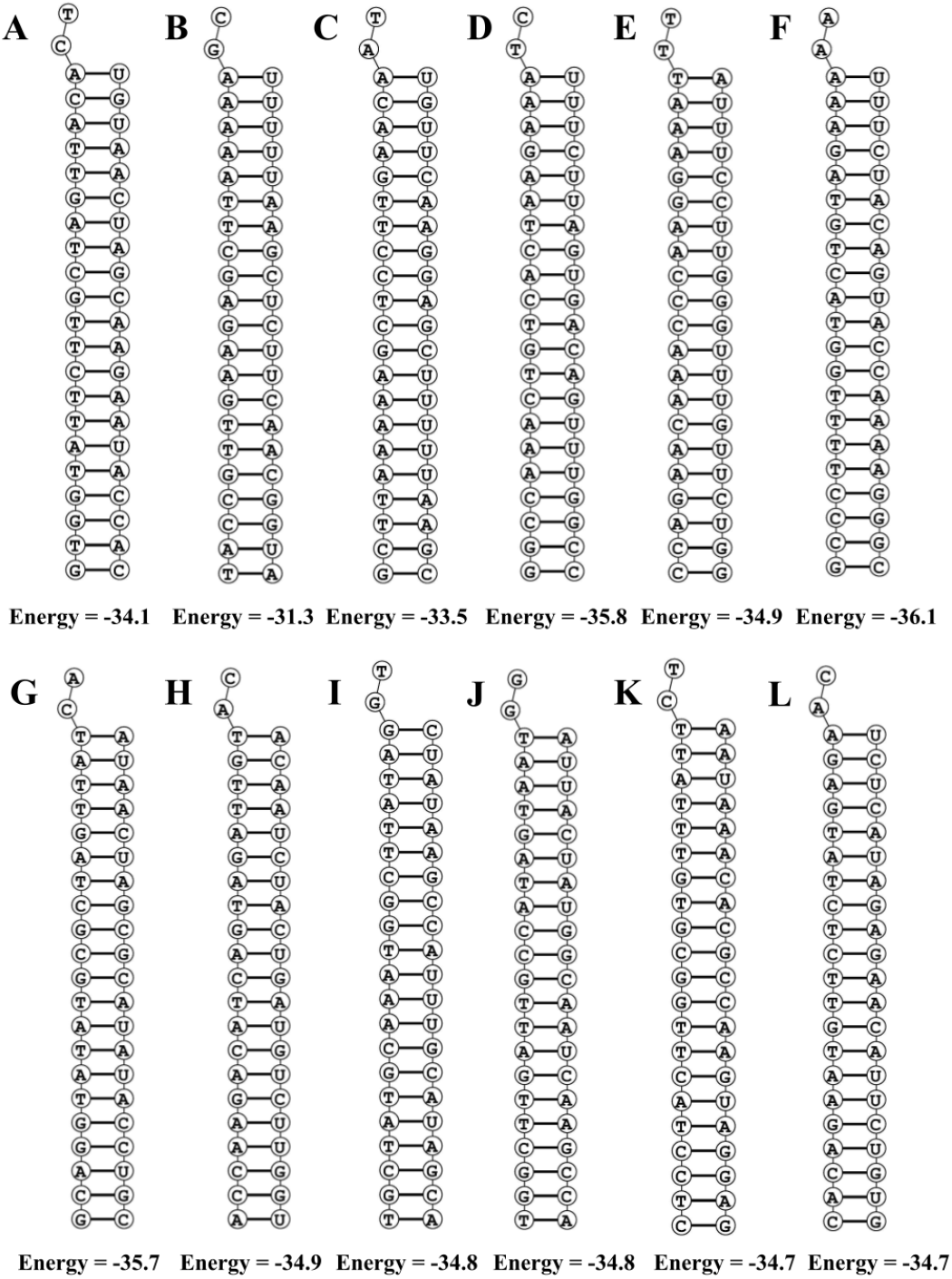
Structure of binding of target RNA and siRNA (guide strand) with corresponding predicted minimum free energy. The structures are for A. e5 B. m1 C. m2 D. n1 E. n2 F. g36 G. g82 H. g83 I. g105 J. g134 K. g80 L. g109 siRNAs.

The three-dimensional structure of Human Ago2 protein (PDB ID: 4F3T) that we considered for the docking studies had 90.7% residues in the core region of the Ramachandran plot thereby qualifying as a good quality structure (Figure 3b). On the other hand, we modelled siRNA using the 3dRNA webserver which builds RNA 3D structures from sequences by using the smallest secondary elements (SSEs)[39].

The Ago2 and modelled siRNAs were docked with the help of the HDOCK server. To perform traditional global docking, the server employs a hybrid-docking technique that combines template-based modelling and free docking, as well as follows a hierarchical FFT-based docking algorithm. Resultant Ago2-siRNAs docked complexes were retrieved from the server and manually analysed to determine the optimal docked complex based on docking score, interaction pattern analysis, and placement of siRNA within the Ago2 domains (PAZ, MID, N-terminal and PIWI domain) (Figure 3a).

Based on the interaction pattern analysis of 12 filtered siRNAs guide strand with target mRNA (Table 3), 4 best final siRNAs (e5, m2, n1 and g80) were sorted and hypothesized for each gene that could mediate the post-transcriptional gene silencing of the targeted gene (Table 4). The docking score of final siRNAs (e5, m2, n1 and g80) showed impressive binding with argonaute protein within the docked complex (Figure 6), ranging from -322.34 to -358.89, where g80 had the highest binding affinity facilitated by the formation of 4 salt bridges, 8 H-bonds, 10 electrostatic bonds and 2 hydrophobic bonds involving ARG255, LYS263, ARG280, ARG286, GLY331, GLU333, ARG351, LYS354, ALA369, ARG710, HIS712, ARG714, ILE756, ARG761, TYR790, ARG792 and TYR804 residues. The detailed elucidation about interacting residues is provided in Supplementary Table 14.

**Table 4.**
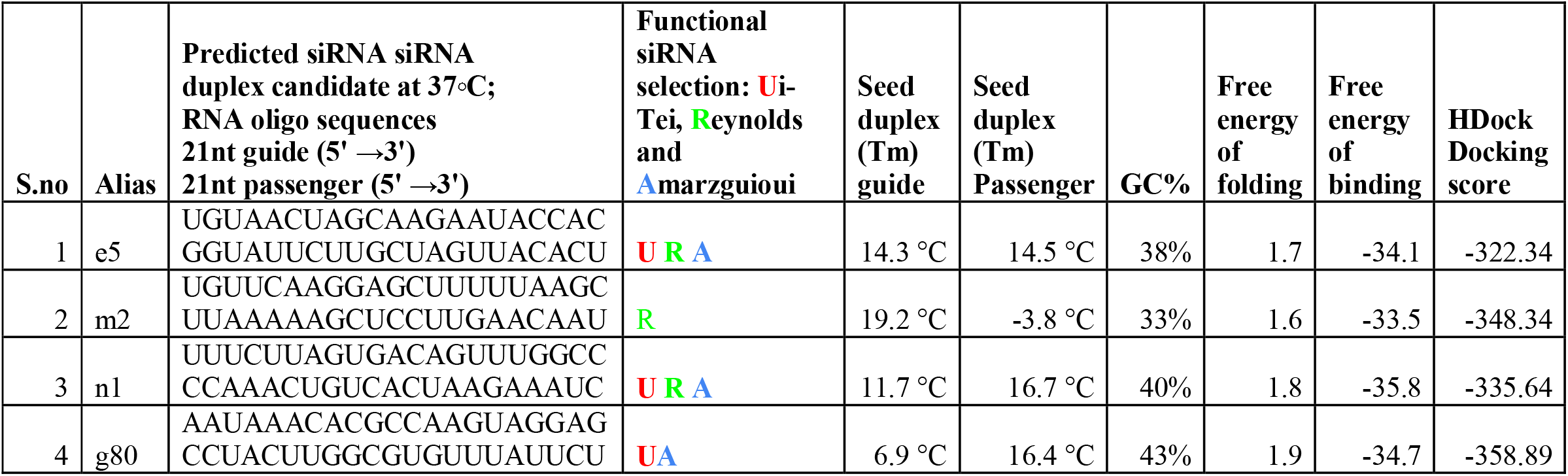
Final siRNA candidates with respect to each gene.

**Figure 6.**
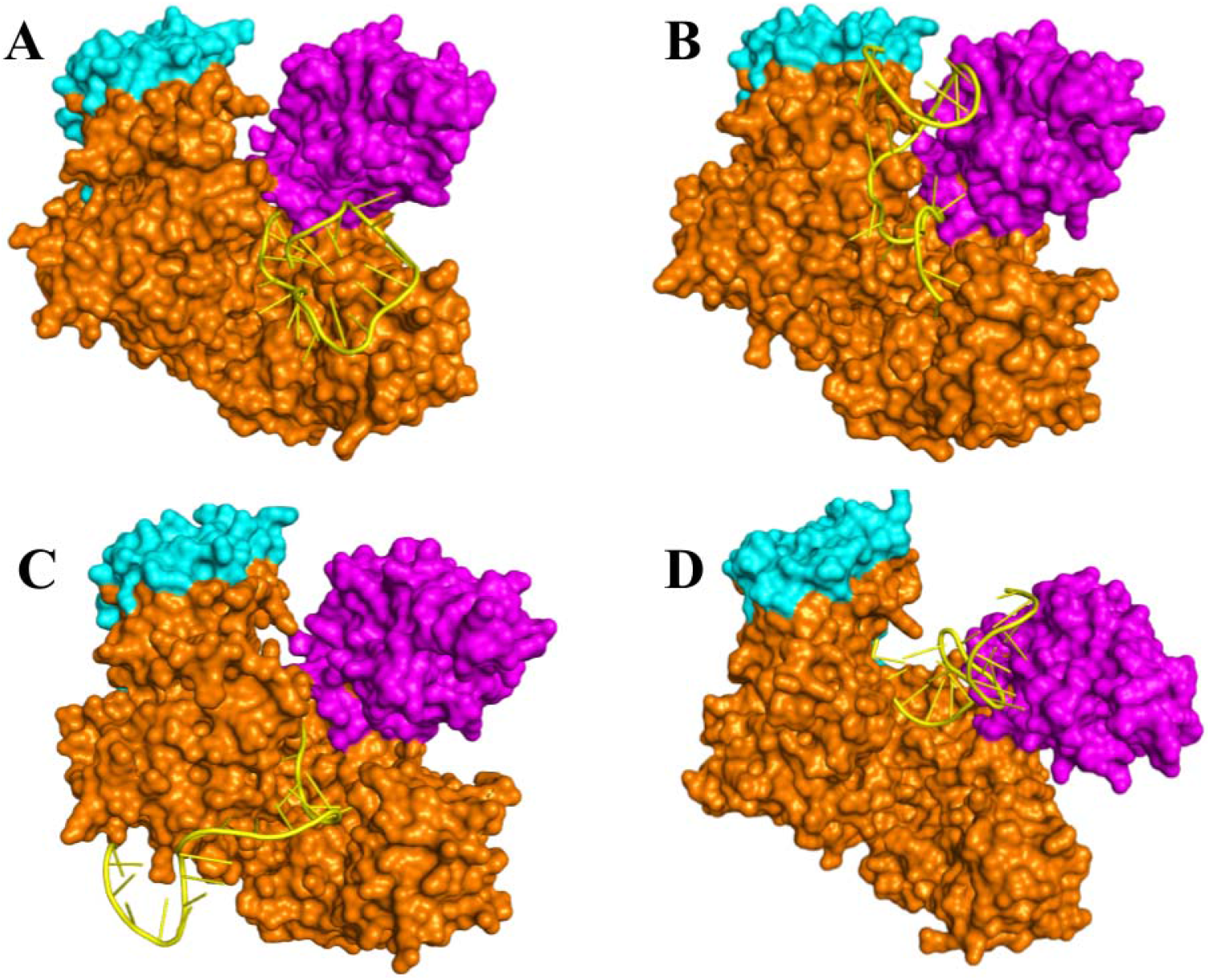
Docked structures of final siRNA candidates (cartoon view) with Human Ago2 protein (surface view). The PAZ domain and the MID domain are coloured as magenta and cyan respectively and rest of the protein denoted by orange colour (including N-terminal domain and PIWI domain). The siRNA was designated by yellow colour. The structures are for A. e5 B. m2 C. n1 D. n1 E. g80 siRNAs.

Therapeutic applications of siRNAs are quite challenging as they have several issues, such as siRNA instability, limited cellular uptake, and the lack of a competent delivery method[49]. However, an appropriate promoter-controlled vector can aid in the targeting of therapeutic genes to the targeted cell for efficient gene therapy[50]. Vector-based siRNA in plasmid form can also be used to target desired genes within a given cell culture to examine the potentiality of a newly created siRNA[51]. Furthermore, plasmids containing siRNA can be directly delivered into the organs of choice[52].

In this present study, four potential siRNA candidates were hypothesized to be effective at binding and cleaving specific SARS-CoV-2 (Indian strains) mRNA targets of structural proteins (Table 4). The suggested therapeutic molecule may be used to treat the COVID-19 pandemic in India on a broad scale because since the study comprises a large array of 811 SARS-CoV-2 sequences from all over India, however, it needs further validation in *in vitro* and *in vivo* studies.

## Conclusions

siRNA therapy might be a promising tool of the RNAi pathway for controlling viral infections in humans by PTGS of the significant gene in various biological systems. In this study, four siRNA molecules were predicted to be effective against the envelope gene (E), membrane gene (M), nucleocapsid phosphoprotein gene (N) and surface glycoprotein gene (S) of 811 Indian strains of the SARS-CoV-2 virus using various computational tools considering all maximum parameters in prominent conditions molecular modelling and docking analysis. However, future applications will necessitate additional validations *in vitro* and *in vivo* animal models. In the battle against the Covid-19 pandemic, these synthetic molecules may be used as novel antiviral therapy and provide a basis for the researchers and pharmaceutical industry to create antiviral therapeutics at the genome level.

## Abbreviations

COVID-19: COrona VIrus Disease 2019
SARS-CoV-2: Severe Acute Respiratory Syndrome-Corona Virus-2
siRNA: Small interfering RNA
PTGS: Post-Transcriptional Gene Silencing
RNAi: RNA interference
Tm: Melting temperature

## CRediT authorship contribution statement

**Premnath Madanagopal:** Conceptualization, Methodology, Data Curation, Visualization, Writing - Original Draft, Writing - Review & Editing. **Harshini Muthukumar:** Visualization, Data Curation. **Kothai Thiruvengadam:** Writing - Review & Editing, Supervision

## Data availability

All necessary data generated or analyzed during this study are included in this article. Any additional data could be available from the corresponding author upon request.

## Acknowledgement

K.T. thank BIC at DoBT, AU (BT/PR40163/BTIS/137/31/2021), Department of Biotechnology, Government of India for computational facilities and manpower support.

