## Supplementary Table 1 for "Computational study and design of effective siRNAs to silence structural proteins associated genes of Indian SARS-CoV-2 strains"

**Supplementary Table 1.** Accession number, location and uploaded date of the SARS-CoV-2 Indian strains and reference Strain used in the Study.

| **S.no** | **Geo Location** | **Nucleotide Accession** | **Date (Uploaded in NCBI)** |
| --- | --- | --- | --- |
| 1 | Asia; China | NC_045512.2 | 18-Jul-20 |
| 2 | Asia; India | MZ292158.1 | 26-May-21 |
| 3 | Asia; India | MZ292157.1 | 26-May-21 |
| 4 | Asia; India | MZ292155.1 | 26-May-21 |
| 5 | Asia; India | MZ292154.1 | 26-May-21 |
| 6 | Asia; India | MZ292153.1 | 26-May-21 |
| 7 | Asia; India | MZ292152.1 | 26-May-21 |
| 8 | Asia; India | MZ292151.1 | 26-May-21 |
| 9 | Asia; India | MZ292150.1 | 26-May-21 |
| 10 | Asia; India | MZ292148.1 | 26-May-21 |
| 11 | Asia; India | MZ292147.1 | 26-May-21 |
| 12 | Asia; India | MZ292146.1 | 26-May-21 |
| 13 | Asia; India | MZ292145.1 | 26-May-21 |
| 14 | Asia; India | MZ292138.1 | 26-May-21 |
| 15 | Asia; India | MZ292137.1 | 26-May-21 |
| 16 | Asia; India | MZ292136.1 | 26-May-21 |
| 17 | Asia; India | MZ292135.1 | 26-May-21 |
| 18 | Asia; India | MZ292134.1 | 26-May-21 |
| 19 | Asia; India | MZ292133.1 | 26-May-21 |
| 20 | Asia; India | MZ292131.1 | 26-May-21 |
| 21 | Asia; India | MZ292130.1 | 26-May-21 |
| 22 | Asia; India | MZ292129.1 | 26-May-21 |
| 23 | Asia; India | MZ292128.1 | 26-May-21 |
| 24 | Asia; India | MZ292127.1 | 26-May-21 |
| 25 | Asia; India | MZ292126.1 | 26-May-21 |
| 26 | Asia; India | MZ292094.1 | 25-May-21 |
| 27 | Asia; India | MW828340.1 | 30-Mar-21 |
| 28 | Asia; India | MW181831.2 | 09-Nov-20 |
| 29 | Asia; India | MW181829.1 | 28-Oct-20 |
| 30 | Asia; India | MT940473.1 | 29-Aug-20 |
| 31 | Asia; India | MT940472.1 | 29-Aug-20 |
| 32 | Asia; India | MT800010.1 | 27-Jul-20 |
| 33 | Asia; India | MT799991.1 | 27-Jul-20 |
| 34 | Asia; India | MT772293.1 | 17-Jul-20 |
| 35 | Asia; India | MT772271.1 | 17-Jul-20 |
| 36 | Asia; India | MT759714.1 | 15-Jul-20 |
| 37 | Asia; India | MT740923.1 | 10-Jul-20 |
| 38 | Asia; India | MT664197.1 | 25-Jun-20 |
| 39 | Asia; India | MT635409.1 | 18-Jun-20 |
| 40 | Asia; India | MT576034.1 | 08-Jun-20 |
| 41 | Asia; India | MT539174.1 | 30-May-20 |
| 42 | Asia; India | MT539170.1 | 30-May-20 |
| 43 | Asia; India | MT509511.1 | 23-May-20 |
| 44 | Asia; India | MT509506.1 | 23-May-20 |
| 45 | Asia; India | MT451877.1 | 11-May-20 |
| 46 | Asia; India | MW828349.1 | 30-Mar-21 |
| 47 | Asia; India | MW828343.1 | 30-Mar-21 |
| 48 | Asia; India | MW828339.1 | 30-Mar-21 |
| 49 | Asia; India | MW828330.1 | 30-Mar-21 |
| 50 | Asia; India | MW559533.2 | 19-Feb-21 |
| 51 | Asia; India | MT050493.1 | 06-Apr-20 |
| 52 | Asia; India | MT012098.1 | 06-Apr-20 |
| 53 | Asia; India | MZ269147.1 | 24-May-21 |
| 54 | Asia; India | MZ268619.1 | 24-May-21 |
| 55 | Asia; India | MZ268601.1 | 24-May-21 |
| 56 | Asia; India | MZ262335.1 | 21-May-21 |
| 57 | Asia; India | MZ262332.1 | 21-May-21 |
| 58 | Asia; India | MZ262306.1 | 21-May-21 |
| 59 | Asia; India | MZ262303.1 | 21-May-21 |
| 60 | Asia; India | MZ262297.1 | 21-May-21 |
| 61 | Asia; India | MZ262295.1 | 21-May-21 |
| 62 | Asia; India | MZ254991.1 | 21-May-21 |
| 63 | Asia; India | MZ203530.1 | 15-May-21 |
| 64 | Asia; India | MW828341.1 | 30-Mar-21 |
| 65 | Asia; India | MW819940.1 | 28-Mar-21 |
| 66 | Asia; India | MW595948.1 | 12-Feb-21 |
| 67 | Asia; India | MW595916.1 | 12-Feb-21 |
| 68 | Asia; India | MW595915.1 | 12-Feb-21 |
| 69 | Asia; India | MW595914.1 | 12-Feb-21 |
| 70 | Asia; India | MW595912.1 | 12-Feb-21 |
| 71 | Asia; India | MW555786.1 | 02-Feb-21 |
| 72 | Asia; India | MW555597.1 | 02-Feb-21 |
| 73 | Asia; India | MW242962.1 | 12-Nov-20 |
| 74 | Asia; India | MW242706.1 | 11-Nov-20 |
| 75 | Asia; India | MW242705.1 | 11-Nov-20 |
| 76 | Asia; India | MW242704.1 | 11-Nov-20 |
| 77 | Asia; India | MW242671.1 | 11-Nov-20 |
| 78 | Asia; India | MW181847.1 | 28-Oct-20 |
| 79 | Asia; India | MW181832.1 | 28-Oct-20 |
| 80 | Asia; India | MW173254.1 | 27-Oct-20 |
| 81 | Asia; India | MW173253.1 | 27-Oct-20 |
| 82 | Asia; India | MW173252.1 | 27-Oct-20 |
| 83 | Asia; India | MW173250.1 | 27-Oct-20 |
| 84 | Asia; India | MW173245.1 | 27-Oct-20 |
| 85 | Asia; India | MW166864.1 | 26-Oct-20 |
| 86 | Asia; India | MW165872.1 | 26-Oct-20 |
| 87 | Asia; India | MW165869.1 | 26-Oct-20 |
| 88 | Asia; India | MT954971.1 | 03-Sep-20 |
| 89 | Asia; India | MT954959.1 | 03-Sep-20 |
| 90 | Asia; India | MT954593.1 | 03-Sep-20 |
| 91 | Asia; India | MT954416.1 | 03-Sep-20 |
| 92 | Asia; India | MT954072.1 | 03-Sep-20 |
| 93 | Asia; India | MT951208.1 | 02-Sep-20 |
| 94 | Asia; India | MT951177.1 | 02-Sep-20 |
| 95 | Asia; India | MT951170.1 | 02-Sep-20 |
| 96 | Asia; India | MT950684.1 | 02-Sep-20 |
| 97 | Asia; India | MT950532.1 | 02-Sep-20 |
| 98 | Asia; India | MT941272.1 | 31-Aug-20 |
| 99 | Asia; India | MT808071.1 | 28-Jul-20 |
| 100 | Asia; India | MT806176.1 | 28-Jul-20 |
| 101 | Asia; India | MT806104.1 | 28-Jul-20 |
| 102 | Asia; India | MT801043.1 | 27-Jul-20 |
| 103 | Asia; India | MT800939.1 | 27-Jul-20 |
| 104 | Asia; India | MT800924.1 | 27-Jul-20 |
| 105 | Asia; India | MT800895.1 | 27-Jul-20 |
| 106 | Asia; India | MT800794.1 | 27-Jul-20 |
| 107 | Asia; India | MT800753.1 | 27-Jul-20 |
| 108 | Asia; India | MT800282.1 | 27-Jul-20 |
| 109 | Asia; India | MT800047.1 | 27-Jul-20 |
| 110 | Asia; India | MT800045.1 | 27-Jul-20 |
| 111 | Asia; India | MT800031.1 | 27-Jul-20 |
| 112 | Asia; India | MT800030.1 | 27-Jul-20 |
| 113 | Asia; India | MT800029.1 | 27-Jul-20 |
| 114 | Asia; India | MT800024.1 | 27-Jul-20 |
| 115 | Asia; India | MT800021.1 | 27-Jul-20 |
| 116 | Asia; India | MT800005.1 | 27-Jul-20 |
| 117 | Asia; India | MT800002.1 | 27-Jul-20 |
| 118 | Asia; India | MT799974.1 | 27-Jul-20 |
| 119 | Asia; India | MT799967.1 | 27-Jul-20 |
| 120 | Asia; India | MT773134.1 | 17-Jul-20 |
| 121 | Asia; India | MT773133.1 | 17-Jul-20 |
| 122 | Asia; India | MT772297.1 | 17-Jul-20 |
| 123 | Asia; India | MT772294.1 | 17-Jul-20 |
| 124 | Asia; India | MT772252.1 | 17-Jul-20 |
| 125 | Asia; India | MT772240.1 | 17-Jul-20 |
| 126 | Asia; India | MT758701.1 | 15-Jul-20 |
| 127 | Asia; India | MT758421.1 | 15-Jul-20 |
| 128 | Asia; India | MT758202.1 | 15-Jul-20 |
| 129 | Asia; India | MT758197.1 | 15-Jul-20 |
| 130 | Asia; India | MT758193.1 | 15-Jul-20 |
| 131 | Asia; India | MT741096.1 | 10-Jul-20 |
| 132 | Asia; India | MT740686.1 | 10-Jul-20 |
| 133 | Asia; India | MT740483.1 | 10-Jul-20 |
| 134 | Asia; India | MT740441.1 | 10-Jul-20 |
| 135 | Asia; India | MT676397.1 | 29-Jun-20 |
| 136 | Asia; India | MT676389.1 | 29-Jun-20 |
| 137 | Asia; India | MT676388.1 | 29-Jun-20 |
| 138 | Asia; India | MT676369.1 | 29-Jun-20 |
| 139 | Asia; India | MT676290.1 | 29-Jun-20 |
| 140 | Asia; India | MT676289.1 | 29-Jun-20 |
| 141 | Asia; India | MT676288.1 | 29-Jun-20 |
| 142 | Asia; India | MT676014.1 | 29-Jun-20 |
| 143 | Asia; India | MT676013.1 | 29-Jun-20 |
| 144 | Asia; India | MT675951.1 | 29-Jun-20 |
| 145 | Asia; India | MT664795.1 | 25-Jun-20 |
| 146 | Asia; India | MT664793.1 | 25-Jun-20 |
| 147 | Asia; India | MT664169.1 | 25-Jun-20 |
| 148 | Asia; India | MT635410.1 | 18-Jun-20 |
| 149 | Asia; India | MT635408.1 | 18-Jun-20 |
| 150 | Asia; India | MT635397.1 | 18-Jun-20 |
| 151 | Asia; India | MT635339.1 | 18-Jun-20 |
| 152 | Asia; India | MT635272.1 | 18-Jun-20 |
| 153 | Asia; India | MT607619.1 | 15-Jun-20 |
| 154 | Asia; India | MT607612.1 | 15-Jun-20 |
| 155 | Asia; India | MT577014.1 | 21-Sep-20 |
| 156 | Asia; India | MT577013.1 | 21-Sep-20 |
| 157 | Asia; India | MT577012.1 | 21-Sep-20 |
| 158 | Asia; India | MT577011.1 | 21-Sep-20 |
| 159 | Asia; India | MT577010.1 | 21-Sep-20 |
| 160 | Asia; India | MT577009.1 | 21-Sep-20 |
| 161 | Asia; India | MT576530.1 | 08-Jun-20 |
| 162 | Asia; India | MT576053.1 | 08-Jun-20 |
| 163 | Asia; India | MT576052.1 | 08-Jun-20 |
| 164 | Asia; India | MT576049.1 | 08-Jun-20 |
| 165 | Asia; India | MT576045.1 | 08-Jun-20 |
| 166 | Asia; India | MT576041.1 | 08-Jun-20 |
| 167 | Asia; India | MT576040.1 | 08-Jun-20 |
| 168 | Asia; India | MT576039.1 | 08-Jun-20 |
| 169 | Asia; India | MT576031.1 | 08-Jun-20 |
| 170 | Asia; India | MT560827.1 | 04-Jun-20 |
| 171 | Asia; India | MT560694.1 | 04-Jun-20 |
| 172 | Asia; India | MT560690.1 | 04-Jun-20 |
| 173 | Asia; India | MT560675.1 | 04-Jun-20 |
| 174 | Asia; India | MT560673.1 | 04-Jun-20 |
| 175 | Asia; India | MT560669.1 | 04-Jun-20 |
| 176 | Asia; India | MT560657.1 | 04-Jun-20 |
| 177 | Asia; India | MT539175.1 | 30-May-20 |
| 178 | Asia; India | MT539173.1 | 30-May-20 |
| 179 | Asia; India | MT539167.1 | 30-May-20 |
| 180 | Asia; India | MT539164.1 | 30-May-20 |
| 181 | Asia; India | MT509659.1 | 24-May-20 |
| 182 | Asia; India | MT509650.1 | 24-May-20 |
| 183 | Asia; India | MT509509.1 | 23-May-20 |
| 184 | Asia; India | MT509495.1 | 23-May-20 |
| 185 | Asia; India | MT509494.1 | 23-May-20 |
| 186 | Asia; India | MT496997.1 | 20-May-20 |
| 187 | Asia; India | MT496996.1 | 20-May-20 |
| 188 | Asia; India | MT496990.1 | 20-May-20 |
| 189 | Asia; India | MT496989.1 | 20-May-20 |
| 190 | Asia; India | MT496988.1 | 20-May-20 |
| 191 | Asia; India | MT496984.1 | 20-May-20 |
| 192 | Asia; India | MT496977.1 | 20-May-20 |
| 193 | Asia; India | MT496976.1 | 20-May-20 |
| 194 | Asia; India | MT496975.1 | 20-May-20 |
| 195 | Asia; India | MT496973.1 | 20-May-20 |
| 196 | Asia; India | MT496972.1 | 20-May-20 |
| 197 | Asia; India | MT483702.1 | 19-May-20 |
| 198 | Asia; India | MT483559.1 | 19-May-20 |
| 199 | Asia; India | MT483557.1 | 19-May-20 |
| 200 | Asia; India | MT483555.1 | 19-May-20 |
| 201 | Asia; India | MT483553.1 | 19-May-20 |
| 202 | Asia; India | MT481909.1 | 20-May-20 |
| 203 | Asia; India | MT481908.1 | 20-May-20 |
| 204 | Asia; India | MT481907.1 | 20-May-20 |
| 205 | Asia; India | MT481905.1 | 20-May-20 |
| 206 | Asia; India | MT481904.1 | 20-May-20 |
| 207 | Asia; India | MT481903.1 | 20-May-20 |
| 208 | Asia; India | MT481902.1 | 20-May-20 |
| 209 | Asia; India | MT481900.1 | 20-May-20 |
| 210 | Asia; India | MT481896.1 | 20-May-20 |
| 211 | Asia; India | MT467260.1 | 14-May-20 |
| 212 | Asia; India | MT467259.1 | 14-May-20 |
| 213 | Asia; India | MT467257.1 | 14-May-20 |
| 214 | Asia; India | MT467255.1 | 14-May-20 |
| 215 | Asia; India | MT467251.1 | 14-May-20 |
| 216 | Asia; India | MT467250.1 | 14-May-20 |
| 217 | Asia; India | MT467249.1 | 14-May-20 |
| 218 | Asia; India | MT467246.1 | 14-May-20 |
| 219 | Asia; India | MT467244.1 | 14-May-20 |
| 220 | Asia; India | MT467242.1 | 14-May-20 |
| 221 | Asia; India | MT467239.1 | 14-May-20 |
| 222 | Asia; India | MT467238.1 | 14-May-20 |
| 223 | Asia; India | MT467237.1 | 14-May-20 |
| 224 | Asia; India | MT457402.1 | 12-May-20 |
| 225 | Asia; India | MT451887.1 | 11-May-20 |
| 226 | Asia; India | MT451886.1 | 11-May-20 |
| 227 | Asia; India | MT451885.1 | 11-May-20 |
| 228 | Asia; India | MT451884.1 | 11-May-20 |
| 229 | Asia; India | MT451883.1 | 11-May-20 |
| 230 | Asia; India | MT451876.1 | 11-May-20 |
| 231 | Asia; India | MT435084.1 | 06-May-20 |
| 232 | Asia; India | MT435083.1 | 06-May-20 |
| 233 | Asia; India | MT435082.1 | 06-May-20 |
| 234 | Asia; India | MT435081.1 | 06-May-20 |
| 235 | Asia; India | MT435080.1 | 06-May-20 |
| 236 | Asia; India | MT435079.1 | 06-May-20 |
| 237 | Asia; India | MT415321.1 | 30-Apr-20 |
| 238 | Asia; India | MZ268620.1 | 24-May-21 |
| 239 | Asia; India | MZ268600.1 | 24-May-21 |
| 240 | Asia; India | MZ261822.1 | 21-May-21 |
| 241 | Asia; India | MZ261819.1 | 21-May-21 |
| 242 | Asia; India | MZ261812.1 | 21-May-21 |
| 243 | Asia; India | MZ227612.1 | 19-May-21 |
| 244 | Asia; India | MZ227582.1 | 19-May-21 |
| 245 | Asia; India | MZ227545.1 | 19-May-21 |
| 246 | Asia; India | MW927136.1 | 15-Apr-21 |
| 247 | Asia; India | MW645476.1 | 22-Feb-21 |
| 248 | Asia; India | MW425563.1 | 01-Jan-21 |
| 249 | Asia; India | MW181846.1 | 28-Oct-20 |
| 250 | Asia; India | MT806191.1 | 28-Jul-20 |
| 251 | Asia; India | MT669321.1 | 26-Jun-20 |
| 252 | Asia; India | MT577015.1 | 21-Sep-20 |
| 253 | Asia; India | MT415323.1 | 30-Apr-20 |
| 254 | Asia; India | MT953890.1 | 03-Sep-20 |
| 255 | Asia; India | MT953887.1 | 03-Sep-20 |
| 256 | Asia; India | MZ269225.1 | 24-May-21 |
| 257 | Asia; India | MZ269146.1 | 24-May-21 |
| 258 | Asia; India | MZ269094.1 | 24-May-21 |
| 259 | Asia; India | MZ267530.1 | 23-May-21 |
| 260 | Asia; India | MZ267529.1 | 23-May-21 |
| 261 | Asia; India | MZ266572.1 | 22-May-21 |
| 262 | Asia; India | MZ266543.1 | 22-May-21 |
| 263 | Asia; India | MT940526.1 | 29-Aug-20 |
| 264 | Asia; India | MZ269224.1 | 24-May-21 |
| 265 | Asia; India | MZ268638.1 | 24-May-21 |
| 266 | Asia; India | MZ268637.1 | 24-May-21 |
| 267 | Asia; India | MZ268635.1 | 24-May-21 |
| 268 | Asia; India | MZ268629.1 | 24-May-21 |
| 269 | Asia; India | MZ268603.1 | 24-May-21 |
| 270 | Asia; India | MZ268602.1 | 24-May-21 |
| 271 | Asia; India | MZ268209.1 | 24-May-21 |
| 272 | Asia; India | MZ261914.1 | 21-May-21 |
| 273 | Asia; India | MZ261895.1 | 21-May-21 |
| 274 | Asia; India | MZ261834.1 | 21-May-21 |
| 275 | Asia; India | MZ203529.1 | 15-May-21 |
| 276 | Asia; India | MW600627.1 | 16-Feb-21 |
| 277 | Asia; India | MW600453.1 | 16-Feb-21 |
| 278 | Asia; India | MW600436.1 | 16-Feb-21 |
| 279 | Asia; India | MW600363.1 | 16-Feb-21 |
| 280 | Asia; India | MW595982.1 | 12-Feb-21 |
| 281 | Asia; India | MW595950.1 | 12-Feb-21 |
| 282 | Asia; India | MW425837.1 | 01-Jan-21 |
| 283 | Asia; India | MW243003.1 | 12-Nov-20 |
| 284 | Asia; India | MW242999.1 | 12-Nov-20 |
| 285 | Asia; India | MW242786.1 | 11-Nov-20 |
| 286 | Asia; India | MW242781.1 | 11-Nov-20 |
| 287 | Asia; India | MW242780.1 | 11-Nov-20 |
| 288 | Asia; India | MW242692.1 | 11-Nov-20 |
| 289 | Asia; India | MW242691.1 | 11-Nov-20 |
| 290 | Asia; India | MW181844.1 | 28-Oct-20 |
| 291 | Asia; India | MT955326.1 | 03-Sep-20 |
| 292 | Asia; India | MT955114.1 | 03-Sep-20 |
| 293 | Asia; India | MT799998.1 | 27-Jul-20 |
| 294 | Asia; India | MT759820.1 | 15-Jul-20 |
| 295 | Asia; India | MT665974.1 | 25-Jun-20 |
| 296 | Asia; India | MT509510.1 | 23-May-20 |
| 297 | Asia; India | MT509507.1 | 23-May-20 |
| 298 | Asia; India | MT416726.2 | 01-Jun-20 |
| 299 | Asia; India | MT416725.2 | 01-Jun-20 |
| 300 | Asia; India | MZ268644.1 | 24-May-21 |
| 301 | Asia; India | MZ268634.1 | 24-May-21 |
| 302 | Asia; India | MZ254839.1 | 21-May-21 |
| 303 | Asia; India | MZ254838.1 | 21-May-21 |
| 304 | Asia; India | MZ254837.1 | 21-May-21 |
| 305 | Asia; India | MZ254787.1 | 21-May-21 |
| 306 | Asia; India | MZ254774.1 | 21-May-21 |
| 307 | Asia; India | MZ254772.1 | 21-May-21 |
| 308 | Asia; India | MZ254771.1 | 21-May-21 |
| 309 | Asia; India | MW881790.1 | 08-Apr-21 |
| 310 | Asia; India | MW600456.1 | 16-Feb-21 |
| 311 | Asia; India | MW600455.1 | 16-Feb-21 |
| 312 | Asia; India | MW596415.1 | 12-Feb-21 |
| 313 | Asia; India | MW596237.1 | 12-Feb-21 |
| 314 | Asia; India | MW595999.1 | 12-Feb-21 |
| 315 | Asia; India | MW595997.1 | 12-Feb-21 |
| 316 | Asia; India | MW595991.1 | 12-Feb-21 |
| 317 | Asia; India | MW595923.1 | 12-Feb-21 |
| 318 | Asia; India | MW595919.1 | 12-Feb-21 |
| 319 | Asia; India | MT806192.1 | 28-Jul-20 |
| 320 | Asia; India | MT800997.1 | 27-Jul-20 |
| 321 | Asia; India | MT800995.1 | 27-Jul-20 |
| 322 | Asia; India | MT800994.1 | 27-Jul-20 |
| 323 | Asia; India | MT800934.1 | 27-Jul-20 |
| 324 | Asia; India | MT800896.1 | 27-Jul-20 |
| 325 | Asia; India | MT800873.1 | 27-Jul-20 |
| 326 | Asia; India | MT800542.1 | 27-Jul-20 |
| 327 | Asia; India | MT800284.1 | 27-Jul-20 |
| 328 | Asia; India | MT772253.1 | 17-Jul-20 |
| 329 | Asia; India | MT772215.1 | 17-Jul-20 |
| 330 | Asia; India | MT759844.1 | 15-Jul-20 |
| 331 | Asia; India | MT759582.1 | 15-Jul-20 |
| 332 | Asia; India | MT758683.1 | 15-Jul-20 |
| 333 | Asia; India | MT758471.1 | 15-Jul-20 |
| 334 | Asia; India | MT758436.1 | 15-Jul-20 |
| 335 | Asia; India | MT758346.1 | 15-Jul-20 |
| 336 | Asia; India | MT758314.1 | 15-Jul-20 |
| 337 | Asia; India | MT758213.1 | 15-Jul-20 |
| 338 | Asia; India | MT741490.1 | 10-Jul-20 |
| 339 | Asia; India | MT740820.1 | 10-Jul-20 |
| 340 | Asia; India | MT740717.1 | 10-Jul-20 |
| 341 | Asia; India | MT740688.1 | 10-Jul-20 |
| 342 | Asia; India | MT740684.1 | 10-Jul-20 |
| 343 | Asia; India | MT740618.1 | 10-Jul-20 |
| 344 | Asia; India | MT676367.1 | 29-Jun-20 |
| 345 | Asia; India | MT676366.1 | 29-Jun-20 |
| 346 | Asia; India | MT676291.1 | 29-Jun-20 |
| 347 | Asia; India | MT675959.1 | 29-Jun-20 |
| 348 | Asia; India | MT675957.1 | 29-Jun-20 |
| 349 | Asia; India | MT675955.1 | 29-Jun-20 |
| 350 | Asia; India | MT675954.1 | 29-Jun-20 |
| 351 | Asia; India | MT675952.1 | 29-Jun-20 |
| 352 | Asia; India | MT675943.1 | 29-Jun-20 |
| 353 | Asia; India | MT675942.1 | 29-Jun-20 |
| 354 | Asia; India | MT675941.1 | 29-Jun-20 |
| 355 | Asia; India | MT675940.1 | 29-Jun-20 |
| 356 | Asia; India | MT675939.1 | 29-Jun-20 |
| 357 | Asia; India | MT675938.1 | 29-Jun-20 |
| 358 | Asia; India | MT675937.1 | 29-Jun-20 |
| 359 | Asia; India | MT675933.1 | 29-Jun-20 |
| 360 | Asia; India | MT665970.1 | 25-Jun-20 |
| 361 | Asia; India | MT665028.1 | 25-Jun-20 |
| 362 | Asia; India | MT665006.1 | 25-Jun-20 |
| 363 | Asia; India | MT664990.1 | 25-Jun-20 |
| 364 | Asia; India | MT664986.1 | 25-Jun-20 |
| 365 | Asia; India | MT664822.1 | 25-Jun-20 |
| 366 | Asia; India | MT664808.1 | 25-Jun-20 |
| 367 | Asia; India | MT664807.1 | 25-Jun-20 |
| 368 | Asia; India | MT664796.1 | 25-Jun-20 |
| 369 | Asia; India | MT664727.1 | 25-Jun-20 |
| 370 | Asia; India | MT664205.1 | 25-Jun-20 |
| 371 | Asia; India | MT664202.1 | 25-Jun-20 |
| 372 | Asia; India | MT664172.1 | 25-Jun-20 |
| 373 | Asia; India | MT664170.1 | 25-Jun-20 |
| 374 | Asia; India | MT664161.1 | 25-Jun-20 |
| 375 | Asia; India | MT664143.1 | 25-Jun-20 |
| 376 | Asia; India | MT635857.1 | 18-Jun-20 |
| 377 | Asia; India | MT635407.1 | 18-Jun-20 |
| 378 | Asia; India | MT635392.1 | 18-Jun-20 |
| 379 | Asia; India | MW242647.1 | 11-Nov-20 |
| 380 | Asia; India | MT801051.1 | 27-Jul-20 |
| 381 | Asia; India | MT801006.1 | 27-Jul-20 |
| 382 | Asia; India | MT800942.1 | 27-Jul-20 |
| 383 | Asia; India | MT800515.1 | 27-Jul-20 |
| 384 | Asia; India | MT800105.1 | 27-Jul-20 |
| 385 | Asia; India | MT800050.1 | 27-Jul-20 |
| 386 | Asia; India | MT800013.1 | 27-Jul-20 |
| 387 | Asia; India | MT740438.1 | 10-Jul-20 |
| 388 | Asia; India | MT635406.1 | 18-Jun-20 |
| 389 | Asia; India | MT635405.1 | 18-Jun-20 |
| 390 | Asia; India | MT635404.1 | 18-Jun-20 |
| 391 | Asia; India | MT635393.1 | 18-Jun-20 |
| 392 | Asia; India | MT607618.1 | 15-Jun-20 |
| 393 | Asia; India | MT576037.1 | 08-Jun-20 |
| 394 | Asia; India | MT576035.1 | 08-Jun-20 |
| 395 | Asia; India | MT539176.1 | 30-May-20 |
| 396 | Asia; India | MT539172.1 | 30-May-20 |
| 397 | Asia; India | MT539171.1 | 30-May-20 |
| 398 | Asia; India | MT539169.1 | 30-May-20 |
| 399 | Asia; India | MT539168.1 | 30-May-20 |
| 400 | Asia; India | MT539166.1 | 30-May-20 |
| 401 | Asia; India | MT539165.1 | 30-May-20 |
| 402 | Asia; India | MT509656.1 | 24-May-20 |
| 403 | Asia; India | MT509649.1 | 24-May-20 |
| 404 | Asia; India | MT950189.1 | 02-Sep-20 |
| 405 | Asia; India | MT799969.1 | 27-Jul-20 |
| 406 | Asia; India | MT758240.1 | 15-Jul-20 |
| 407 | Asia; India | MT666042.1 | 25-Jun-20 |
| 408 | Asia; India | MT509504.1 | 23-May-20 |
| 409 | Asia; India | MT467252.1 | 14-May-20 |
| 410 | Asia; India | MT467247.1 | 14-May-20 |
| 411 | Asia; India | MT467241.1 | 14-May-20 |
| 412 | Asia; India | MT467240.1 | 14-May-20 |
| 413 | Asia; India | MT435086.1 | 06-May-20 |
| 414 | Asia; India | MZ268643.1 | 24-May-21 |
| 415 | Asia; India | MW828336.1 | 30-Mar-21 |
| 416 | Asia; India | MW243586.1 | 12-Nov-20 |
| 417 | Asia; India | MW242779.1 | 11-Nov-20 |
| 418 | Asia; India | MW242689.1 | 11-Nov-20 |
| 419 | Asia; India | MW173463.1 | 27-Oct-20 |
| 420 | Asia; India | MW167112.1 | 26-Oct-20 |
| 421 | Asia; India | MW166214.1 | 26-Oct-20 |
| 422 | Asia; India | MW165876.1 | 26-Oct-20 |
| 423 | Asia; India | MW165862.1 | 26-Oct-20 |
| 424 | Asia; India | MW165857.1 | 26-Oct-20 |
| 425 | Asia; India | MT953991.1 | 03-Sep-20 |
| 426 | Asia; India | MT953981.1 | 03-Sep-20 |
| 427 | Asia; India | MT953893.1 | 03-Sep-20 |
| 428 | Asia; India | MT358637.1 | 20-Apr-20 |
| 429 | Asia; India | MZ262293.1 | 21-May-21 |
| 430 | Asia; India | MZ261927.1 | 21-May-21 |
| 431 | Asia; India | MW191501.1 | 30-Oct-20 |
| 432 | Asia; India | MW181848.1 | 28-Oct-20 |
| 433 | Asia; India | MW181843.1 | 28-Oct-20 |
| 434 | Asia; India | MW181807.1 | 28-Oct-20 |
| 435 | Asia; India | MW181806.1 | 28-Oct-20 |
| 436 | Asia; India | MW181805.1 | 28-Oct-20 |
| 437 | Asia; India | MW181804.1 | 28-Oct-20 |
| 438 | Asia; India | MT950348.1 | 02-Sep-20 |
| 439 | Asia; India | MT889692.1 | 16-Aug-20 |
| 440 | Asia; India | MT758439.1 | 15-Jul-20 |
| 441 | Asia; India | MT664774.1 | 25-Jun-20 |
| 442 | Asia; India | MT635856.1 | 18-Jun-20 |
| 443 | Asia; India | MT635403.1 | 18-Jun-20 |
| 444 | Asia; India | MT635391.1 | 18-Jun-20 |
| 445 | Asia; India | MT635328.1 | 18-Jun-20 |
| 446 | Asia; India | MT635271.1 | 18-Jun-20 |
| 447 | Asia; India | MT635269.1 | 18-Jun-20 |
| 448 | Asia; India | MT608648.1 | 15-Jun-20 |
| 449 | Asia; India | MT607621.1 | 15-Jun-20 |
| 450 | Asia; India | MT607620.1 | 15-Jun-20 |
| 451 | Asia; India | MT607616.1 | 15-Jun-20 |
| 452 | Asia; India | MT607615.1 | 15-Jun-20 |
| 453 | Asia; India | MT607613.1 | 15-Jun-20 |
| 454 | Asia; India | MT607610.1 | 15-Jun-20 |
| 455 | Asia; India | MT607607.1 | 15-Jun-20 |
| 456 | Asia; India | MT607606.1 | 15-Jun-20 |
| 457 | Asia; India | MT607604.1 | 15-Jun-20 |
| 458 | Asia; India | MT607603.1 | 15-Jun-20 |
| 459 | Asia; India | MT607601.1 | 15-Jun-20 |
| 460 | Asia; India | MT607251.1 | 15-Jun-20 |
| 461 | Asia; India | MT607250.1 | 15-Jun-20 |
| 462 | Asia; India | MT607249.1 | 15-Jun-20 |
| 463 | Asia; India | MT607244.1 | 15-Jun-20 |
| 464 | Asia; India | MT576059.1 | 08-Jun-20 |
| 465 | Asia; India | MT576047.1 | 08-Jun-20 |
| 466 | Asia; India | MT576046.1 | 08-Jun-20 |
| 467 | Asia; India | MT576042.1 | 08-Jun-20 |
| 468 | Asia; India | MT576038.1 | 08-Jun-20 |
| 469 | Asia; India | MT576032.1 | 08-Jun-20 |
| 470 | Asia; India | MT560677.1 | 04-Jun-20 |
| 471 | Asia; India | MT509503.1 | 23-May-20 |
| 472 | Asia; India | MT509501.1 | 23-May-20 |
| 473 | Asia; India | MT965767.1 | 04-Sep-20 |
| 474 | Asia; India | MT940479.1 | 29-Aug-20 |
| 475 | Asia; India | MT940477.1 | 29-Aug-20 |
| 476 | Asia; India | MT801048.1 | 27-Jul-20 |
| 477 | Asia; India | MT800977.1 | 27-Jul-20 |
| 478 | Asia; India | MT800940.1 | 27-Jul-20 |
| 479 | Asia; India | MT800928.1 | 27-Jul-20 |
| 480 | Asia; India | MT800816.1 | 27-Jul-20 |
| 481 | Asia; India | MT800758.1 | 27-Jul-20 |
| 482 | Asia; India | MT800043.1 | 27-Jul-20 |
| 483 | Asia; India | MT800041.1 | 27-Jul-20 |
| 484 | Asia; India | MT800036.1 | 27-Jul-20 |
| 485 | Asia; India | MT772256.1 | 17-Jul-20 |
| 486 | Asia; India | MT772237.1 | 17-Jul-20 |
| 487 | Asia; India | MT760790.1 | 15-Jul-20 |
| 488 | Asia; India | MT759742.1 | 15-Jul-20 |
| 489 | Asia; India | MT759588.1 | 15-Jul-20 |
| 490 | Asia; India | MT758402.1 | 15-Jul-20 |
| 491 | Asia; India | MT758203.1 | 15-Jul-20 |
| 492 | Asia; India | MT740896.1 | 10-Jul-20 |
| 493 | Asia; India | MT676390.1 | 29-Jun-20 |
| 494 | Asia; India | MT669322.1 | 26-Jun-20 |
| 495 | Asia; India | MT665972.1 | 25-Jun-20 |
| 496 | Asia; India | MT664729.1 | 25-Jun-20 |
| 497 | Asia; India | MT664209.1 | 25-Jun-20 |
| 498 | Asia; India | MT664203.1 | 25-Jun-20 |
| 499 | Asia; India | MT664201.1 | 25-Jun-20 |
| 500 | Asia; India | MT664118.1 | 25-Jun-20 |
| 501 | Asia; India | MT483560.1 | 19-May-20 |
| 502 | Asia; India | MT483558.1 | 19-May-20 |
| 503 | Asia; India | MT483556.1 | 19-May-20 |
| 504 | Asia; India | MT483554.1 | 19-May-20 |
| 505 | Asia; India | MT481899.1 | 20-May-20 |
| 506 | Asia; India | MT481897.1 | 20-May-20 |
| 507 | Asia; India | MT467262.1 | 14-May-20 |
| 508 | Asia; India | MT467261.1 | 14-May-20 |
| 509 | Asia; India | MT560706.1 | 04-Jun-20 |
| 510 | Asia; India | MZ269242.1 | 24-May-21 |
| 511 | Asia; India | MZ269100.1 | 24-May-21 |
| 512 | Asia; India | MZ269095.1 | 24-May-21 |
| 513 | Asia; India | MZ269092.1 | 24-May-21 |
| 514 | Asia; India | MZ268819.1 | 24-May-21 |
| 515 | Asia; India | MZ268642.1 | 24-May-21 |
| 516 | Asia; India | MZ268641.1 | 24-May-21 |
| 517 | Asia; India | MZ268631.1 | 24-May-21 |
| 518 | Asia; India | MZ268630.1 | 24-May-21 |
| 519 | Asia; India | MZ268627.1 | 24-May-21 |
| 520 | Asia; India | MZ268626.1 | 24-May-21 |
| 521 | Asia; India | MZ268599.1 | 24-May-21 |
| 522 | Asia; India | MZ268208.1 | 24-May-21 |
| 523 | Asia; India | MZ267528.1 | 23-May-21 |
| 524 | Asia; India | MZ266569.1 | 22-May-21 |
| 525 | Asia; India | MZ266567.1 | 22-May-21 |
| 526 | Asia; India | MZ262334.1 | 21-May-21 |
| 527 | Asia; India | MZ262333.1 | 21-May-21 |
| 528 | Asia; India | MZ262304.1 | 21-May-21 |
| 529 | Asia; India | MZ262301.1 | 21-May-21 |
| 530 | Asia; India | MZ261729.1 | 21-May-21 |
| 531 | Asia; India | MZ261728.1 | 21-May-21 |
| 532 | Asia; India | MZ261726.1 | 21-May-21 |
| 533 | Asia; India | MZ261635.1 | 21-May-21 |
| 534 | Asia; India | MZ256065.1 | 21-May-21 |
| 535 | Asia; India | MZ255008.1 | 21-May-21 |
| 536 | Asia; India | MZ255007.1 | 21-May-21 |
| 537 | Asia; India | MZ254992.1 | 21-May-21 |
| 538 | Asia; India | MZ254987.1 | 21-May-21 |
| 539 | Asia; India | MZ254908.1 | 21-May-21 |
| 540 | Asia; India | MZ254903.1 | 21-May-21 |
| 541 | Asia; India | MZ254902.1 | 21-May-21 |
| 542 | Asia; India | MZ254875.1 | 21-May-21 |
| 543 | Asia; India | MZ254770.1 | 21-May-21 |
| 544 | Asia; India | MW595918.1 | 12-Feb-21 |
| 545 | Asia; India | MW595911.1 | 12-Feb-21 |
| 546 | Asia; India | MW590723.1 | 11-Feb-21 |
| 547 | Asia; India | MW590718.1 | 11-Feb-21 |
| 548 | Asia; India | MW555576.1 | 02-Feb-21 |
| 549 | Asia; India | MW555320.1 | 02-Feb-21 |
| 550 | Asia; India | MT951393.1 | 02-Sep-20 |
| 551 | Asia; India | MT940475.1 | 29-Aug-20 |
| 552 | Asia; India | MT940474.1 | 29-Aug-20 |
| 553 | Asia; India | MT576054.1 | 08-Jun-20 |
| 554 | Asia; India | MT576033.1 | 08-Jun-20 |
| 555 | Asia; India | MT560704.1 | 04-Jun-20 |
| 556 | Asia; India | MT560693.1 | 04-Jun-20 |
| 557 | Asia; India | MT560682.1 | 04-Jun-20 |
| 558 | Asia; India | MT953978.1 | 03-Sep-20 |
| 559 | Asia; India | MT800015.1 | 27-Jul-20 |
| 560 | Asia; India | MZ254899.1 | 21-May-21 |
| 561 | Asia; India | MZ254877.1 | 21-May-21 |
| 562 | Asia; India | MZ269093.1 | 24-May-21 |
| 563 | Asia; India | MZ269079.1 | 24-May-21 |
| 564 | Asia; India | MZ269074.1 | 24-May-21 |
| 565 | Asia; India | MZ269072.1 | 24-May-21 |
| 566 | Asia; India | MZ268850.1 | 24-May-21 |
| 567 | Asia; India | MZ266568.1 | 22-May-21 |
| 568 | Asia; India | MZ266565.1 | 22-May-21 |
| 569 | Asia; India | MZ266554.1 | 22-May-21 |
| 570 | Asia; India | MZ266553.1 | 22-May-21 |
| 571 | Asia; India | MZ266551.1 | 22-May-21 |
| 572 | Asia; India | MZ266547.1 | 22-May-21 |
| 573 | Asia; India | MZ262322.1 | 21-May-21 |
| 574 | Asia; India | MZ262298.1 | 21-May-21 |
| 575 | Asia; India | MZ262296.1 | 21-May-21 |
| 576 | Asia; India | MZ262291.1 | 21-May-21 |
| 577 | Asia; India | MZ261734.1 | 21-May-21 |
| 578 | Asia; India | MZ261699.1 | 21-May-21 |
| 579 | Asia; India | MZ255058.1 | 21-May-21 |
| 580 | Asia; India | MZ254905.1 | 21-May-21 |
| 581 | Asia; India | MZ254876.1 | 21-May-21 |
| 582 | Asia; India | MW828337.1 | 30-Mar-21 |
| 583 | Asia; India | MW828326.1 | 30-Mar-21 |
| 584 | Asia; India | MW828325.1 | 30-Mar-21 |
| 585 | Asia; India | MW819642.1 | 27-Mar-21 |
| 586 | Asia; India | MW819619.1 | 27-Mar-21 |
| 587 | Asia; India | MW812293.1 | 26-Mar-21 |
| 588 | Asia; India | MW600654.1 | 16-Feb-21 |
| 589 | Asia; India | MW600463.1 | 16-Feb-21 |
| 590 | Asia; India | MW600461.1 | 16-Feb-21 |
| 591 | Asia; India | MW368700.1 | 16-Dec-20 |
| 592 | Asia; India | MW243587.1 | 12-Nov-20 |
| 593 | Asia; India | MW242965.1 | 12-Nov-20 |
| 594 | Asia; India | MW242964.1 | 12-Nov-20 |
| 595 | Asia; India | MW242963.1 | 12-Nov-20 |
| 596 | Asia; India | MW242961.1 | 12-Nov-20 |
| 597 | Asia; India | MW242960.1 | 12-Nov-20 |
| 598 | Asia; India | MW242950.1 | 12-Nov-20 |
| 599 | Asia; India | MW242778.1 | 11-Nov-20 |
| 600 | Asia; India | MW242670.1 | 11-Nov-20 |
| 601 | Asia; India | MW242666.1 | 11-Nov-20 |
| 602 | Asia; India | MW242663.1 | 11-Nov-20 |
| 603 | Asia; India | MW242653.1 | 11-Nov-20 |
| 604 | Asia; India | MW173460.1 | 27-Oct-20 |
| 605 | Asia; India | MW173298.1 | 27-Oct-20 |
| 606 | Asia; India | MW167078.1 | 26-Oct-20 |
| 607 | Asia; India | MW166349.1 | 26-Oct-20 |
| 608 | Asia; India | MW166221.1 | 26-Oct-20 |
| 609 | Asia; India | MW165885.1 | 26-Oct-20 |
| 610 | Asia; India | MW165873.1 | 26-Oct-20 |
| 611 | Asia; India | MW165865.1 | 26-Oct-20 |
| 612 | Asia; India | MW165861.1 | 26-Oct-20 |
| 613 | Asia; India | MW165854.1 | 26-Oct-20 |
| 614 | Asia; India | MT965769.1 | 04-Sep-20 |
| 615 | Asia; India | MT955354.1 | 03-Sep-20 |
| 616 | Asia; India | MT955347.1 | 03-Sep-20 |
| 617 | Asia; India | MT954414.1 | 03-Sep-20 |
| 618 | Asia; India | MT954103.1 | 03-Sep-20 |
| 619 | Asia; India | MT954071.1 | 03-Sep-20 |
| 620 | Asia; India | MT954065.1 | 03-Sep-20 |
| 621 | Asia; India | MT954060.1 | 03-Sep-20 |
| 622 | Asia; India | MT953996.1 | 03-Sep-20 |
| 623 | Asia; India | MT953984.1 | 03-Sep-20 |
| 624 | Asia; India | MT953980.1 | 03-Sep-20 |
| 625 | Asia; India | MT953925.1 | 03-Sep-20 |
| 626 | Asia; India | MT953895.1 | 03-Sep-20 |
| 627 | Asia; India | MT953894.1 | 03-Sep-20 |
| 628 | Asia; India | MT953891.1 | 03-Sep-20 |
| 629 | Asia; India | MT953877.1 | 03-Sep-20 |
| 630 | Asia; India | MT953876.1 | 03-Sep-20 |
| 631 | Asia; India | MT953874.1 | 03-Sep-20 |
| 632 | Asia; India | MT953868.1 | 03-Sep-20 |
| 633 | Asia; India | MT951392.1 | 02-Sep-20 |
| 634 | Asia; India | MT951181.1 | 02-Sep-20 |
| 635 | Asia; India | MT951157.1 | 02-Sep-20 |
| 636 | Asia; India | MT950761.1 | 02-Sep-20 |
| 637 | Asia; India | MT950725.1 | 02-Sep-20 |
| 638 | Asia; India | MT950349.1 | 02-Sep-20 |
| 639 | Asia; India | MT950231.1 | 02-Sep-20 |
| 640 | Asia; India | MT950230.1 | 02-Sep-20 |
| 641 | Asia; India | MT950195.1 | 02-Sep-20 |
| 642 | Asia; India | MT950193.1 | 02-Sep-20 |
| 643 | Asia; India | MT950192.1 | 02-Sep-20 |
| 644 | Asia; India | MT950191.1 | 02-Sep-20 |
| 645 | Asia; India | MT950190.1 | 02-Sep-20 |
| 646 | Asia; India | MT950188.1 | 02-Sep-20 |
| 647 | Asia; India | MT950187.1 | 02-Sep-20 |
| 648 | Asia; India | MT941403.1 | 31-Aug-20 |
| 649 | Asia; India | MT941276.1 | 31-Aug-20 |
| 650 | Asia; India | MT941274.1 | 31-Aug-20 |
| 651 | Asia; India | MT941067.1 | 31-Aug-20 |
| 652 | Asia; India | MT941061.1 | 31-Aug-20 |
| 653 | Asia; India | MT941060.1 | 31-Aug-20 |
| 654 | Asia; India | MT941059.1 | 31-Aug-20 |
| 655 | Asia; India | MT941056.1 | 31-Aug-20 |
| 656 | Asia; India | MT941055.1 | 31-Aug-20 |
| 657 | Asia; India | MT941052.1 | 31-Aug-20 |
| 658 | Asia; India | MT941047.1 | 31-Aug-20 |
| 659 | Asia; India | MT940478.1 | 29-Aug-20 |
| 660 | Asia; India | MT940476.1 | 29-Aug-20 |
| 661 | Asia; India | MT940464.1 | 29-Aug-20 |
| 662 | Asia; India | MT940462.1 | 29-Aug-20 |
| 663 | Asia; India | MT940461.1 | 29-Aug-20 |
| 664 | Asia; India | MT940460.1 | 29-Aug-20 |
| 665 | Asia; India | MT940459.1 | 29-Aug-20 |
| 666 | Asia; India | MT940458.1 | 29-Aug-20 |
| 667 | Asia; India | MT940452.1 | 29-Aug-20 |
| 668 | Asia; India | MT940451.1 | 29-Aug-20 |
| 669 | Asia; India | MT940449.1 | 29-Aug-20 |
| 670 | Asia; India | MT801004.1 | 27-Jul-20 |
| 671 | Asia; India | MT800999.1 | 27-Jul-20 |
| 672 | Asia; India | MT800978.1 | 27-Jul-20 |
| 673 | Asia; India | MT800936.1 | 27-Jul-20 |
| 674 | Asia; India | MT800925.1 | 27-Jul-20 |
| 675 | Asia; India | MT800923.1 | 27-Jul-20 |
| 676 | Asia; India | MT800845.1 | 27-Jul-20 |
| 677 | Asia; India | MT800820.1 | 27-Jul-20 |
| 678 | Asia; India | MT800818.1 | 27-Jul-20 |
| 679 | Asia; India | MT800505.1 | 27-Jul-20 |
| 680 | Asia; India | MT799976.1 | 27-Jul-20 |
| 681 | Asia; India | MT799972.1 | 27-Jul-20 |
| 682 | Asia; India | MT799970.1 | 27-Jul-20 |
| 683 | Asia; India | MT772210.1 | 17-Jul-20 |
| 684 | Asia; India | MT758375.1 | 15-Jul-20 |
| 685 | Asia; India | MT758215.1 | 15-Jul-20 |
| 686 | Asia; India | MT740724.1 | 10-Jul-20 |
| 687 | Asia; India | MT740723.1 | 10-Jul-20 |
| 688 | Asia; India | MT740708.1 | 10-Jul-20 |
| 689 | Asia; India | MT740706.1 | 10-Jul-20 |
| 690 | Asia; India | MT740619.1 | 10-Jul-20 |
| 691 | Asia; India | MT740617.1 | 10-Jul-20 |
| 692 | Asia; India | MT740485.1 | 10-Jul-20 |
| 693 | Asia; India | MT740444.1 | 10-Jul-20 |
| 694 | Asia; India | MT676368.1 | 29-Jun-20 |
| 695 | Asia; India | MT675950.1 | 29-Jun-20 |
| 696 | Asia; India | MT675945.1 | 29-Jun-20 |
| 697 | Asia; India | MT675944.1 | 29-Jun-20 |
| 698 | Asia; India | MT664117.1 | 25-Jun-20 |
| 699 | Asia; India | MT635858.1 | 18-Jun-20 |
| 700 | Asia; India | MT635855.1 | 18-Jun-20 |
| 701 | Asia; India | MT635270.1 | 18-Jun-20 |
| 702 | Asia; India | MT607617.1 | 15-Jun-20 |
| 703 | Asia; India | MT607614.1 | 15-Jun-20 |
| 704 | Asia; India | MT607611.1 | 15-Jun-20 |
| 705 | Asia; India | MT607609.1 | 15-Jun-20 |
| 706 | Asia; India | MT607608.1 | 15-Jun-20 |
| 707 | Asia; India | MT607605.1 | 15-Jun-20 |
| 708 | Asia; India | MT607602.1 | 15-Jun-20 |
| 709 | Asia; India | MT594112.1 | 11-Jun-20 |
| 710 | Asia; India | MT576532.1 | 08-Jun-20 |
| 711 | Asia; India | MT576531.1 | 08-Jun-20 |
| 712 | Asia; India | MT576529.1 | 08-Jun-20 |
| 713 | Asia; India | MT576061.1 | 08-Jun-20 |
| 714 | Asia; India | MT576060.1 | 08-Jun-20 |
| 715 | Asia; India | MT576058.1 | 08-Jun-20 |
| 716 | Asia; India | MT576057.1 | 08-Jun-20 |
| 717 | Asia; India | MT576056.1 | 08-Jun-20 |
| 718 | Asia; India | MT576055.1 | 08-Jun-20 |
| 719 | Asia; India | MT576048.1 | 08-Jun-20 |
| 720 | Asia; India | MT576044.1 | 08-Jun-20 |
| 721 | Asia; India | MT576043.1 | 08-Jun-20 |
| 722 | Asia; India | MT560692.1 | 04-Jun-20 |
| 723 | Asia; India | MT560689.1 | 04-Jun-20 |
| 724 | Asia; India | MT560680.1 | 04-Jun-20 |
| 725 | Asia; India | MT560679.1 | 04-Jun-20 |
| 726 | Asia; India | MT560674.1 | 04-Jun-20 |
| 727 | Asia; India | MT560671.1 | 04-Jun-20 |
| 728 | Asia; India | MT560670.1 | 04-Jun-20 |
| 729 | Asia; India | MT560668.1 | 04-Jun-20 |
| 730 | Asia; India | MT560667.1 | 04-Jun-20 |
| 731 | Asia; India | MT560656.1 | 04-Jun-20 |
| 732 | Asia; India | MT509651.1 | 24-May-20 |
| 733 | Asia; India | MT509508.1 | 23-May-20 |
| 734 | Asia; India | MT509505.1 | 23-May-20 |
| 735 | Asia; India | MT509502.1 | 23-May-20 |
| 736 | Asia; India | MT509499.1 | 23-May-20 |
| 737 | Asia; India | MT509498.1 | 23-May-20 |
| 738 | Asia; India | MT496995.1 | 20-May-20 |
| 739 | Asia; India | MT496994.1 | 20-May-20 |
| 740 | Asia; India | MT496993.1 | 20-May-20 |
| 741 | Asia; India | MT496992.1 | 20-May-20 |
| 742 | Asia; India | MT496991.1 | 20-May-20 |
| 743 | Asia; India | MT496987.1 | 20-May-20 |
| 744 | Asia; India | MT496986.1 | 20-May-20 |
| 745 | Asia; India | MT496985.1 | 20-May-20 |
| 746 | Asia; India | MT496983.1 | 20-May-20 |
| 747 | Asia; India | MT496982.1 | 20-May-20 |
| 748 | Asia; India | MT496981.1 | 20-May-20 |
| 749 | Asia; India | MT496980.1 | 20-May-20 |
| 750 | Asia; India | MT496974.1 | 20-May-20 |
| 751 | Asia; India | MT481906.1 | 20-May-20 |
| 752 | Asia; India | MT481901.1 | 20-May-20 |
| 753 | Asia; India | MT481898.1 | 20-May-20 |
| 754 | Asia; India | MT481895.1 | 20-May-20 |
| 755 | Asia; India | MT467263.1 | 14-May-20 |
| 756 | Asia; India | MT467258.1 | 14-May-20 |
| 757 | Asia; India | MT467256.1 | 14-May-20 |
| 758 | Asia; India | MT467254.1 | 14-May-20 |
| 759 | Asia; India | MT467253.1 | 14-May-20 |
| 760 | Asia; India | MT467248.1 | 14-May-20 |
| 761 | Asia; India | MT467245.1 | 14-May-20 |
| 762 | Asia; India | MT467243.1 | 14-May-20 |
| 763 | Asia; India | MT451888.1 | 11-May-20 |
| 764 | Asia; India | MT451882.1 | 11-May-20 |
| 765 | Asia; India | MT451881.1 | 11-May-20 |
| 766 | Asia; India | MT435085.1 | 06-May-20 |
| 767 | Asia; India | MZ262323.1 | 21-May-21 |
| 768 | Asia; India | MZ269101.1 | 24-May-21 |
| 769 | Asia; India | MT560705.1 | 04-Jun-20 |
| 770 | Asia; India | MT954069.1 | 03-Sep-20 |
| 771 | Asia; India | MT954067.1 | 03-Sep-20 |
| 772 | Asia; India | MT740436.1 | 10-Jul-20 |
| 773 | Asia; India | MT496979.1 | 20-May-20 |
| 774 | Asia; India | MT953875.1 | 03-Sep-20 |
| 775 | Asia; India | MT560672.1 | 04-Jun-20 |
| 776 | Asia; India | MW242708.1 | 11-Nov-20 |
| 777 | Asia; India | MZ269263.1 | 24-May-21 |
| 778 | Asia; India | MW193967.2 | 09-Nov-20 |
| 779 | Asia; India | MW191512.1 | 30-Oct-20 |
| 780 | Asia; India | MW191511.1 | 30-Oct-20 |
| 781 | Asia; India | MW191510.1 | 30-Oct-20 |
| 782 | Asia; India | MW191509.1 | 30-Oct-20 |
| 783 | Asia; India | MW181828.1 | 28-Oct-20 |
| 784 | Asia; India | MT415320.1 | 30-Apr-20 |
| 785 | Asia; India | MW555317.1 | 02-Feb-21 |
| 786 | Asia; India | MW555280.1 | 02-Feb-21 |
| 787 | Asia; India | MW422884.1 | 31-Dec-20 |
| 788 | Asia; India | MW181840.1 | 28-Oct-20 |
| 789 | Asia; India | MW181839.1 | 28-Oct-20 |
| 790 | Asia; India | MW181838.2 | 09-Nov-20 |
| 791 | Asia; India | MW181837.1 | 28-Oct-20 |
| 792 | Asia; India | MW181835.1 | 28-Oct-20 |
| 793 | Asia; India | MW181834.1 | 28-Oct-20 |
| 794 | Asia; India | MW181833.1 | 28-Oct-20 |
| 795 | Asia; India | MW181811.1 | 28-Oct-20 |
| 796 | Asia; India | MW181803.1 | 28-Oct-20 |
| 797 | Asia; India | MW181797.1 | 28-Oct-20 |
| 798 | Asia; India | MT509959.1 | 25-May-20 |
| 799 | Asia; India | MT509496.1 | 23-May-20 |
| 800 | Asia; India | MT477885.1 | 18-May-20 |
| 801 | Asia; India | MT457403.1 | 12-May-20 |
| 802 | Asia; India | MT451889.1 | 11-May-20 |
| 803 | Asia; India | MT451880.1 | 11-May-20 |
| 804 | Asia; India | MT451878.1 | 11-May-20 |
| 805 | Asia; India | MT415322.1 | 30-Apr-20 |
| 806 | Asia; India | MW181845.1 | 28-Oct-20 |
| 807 | Asia; India | MT509658.1 | 24-May-20 |
| 808 | Asia; India | MT509657.1 | 24-May-20 |
| 809 | Asia; India | MT509512.1 | 23-May-20 |
| 810 | Asia; India | MT509500.1 | 23-May-20 |
| 811 | Asia; India | MT509497.1 | 23-May-20 |
| 812 | Asia; India | MT451874.1 | 11-May-20 |
