## Supplementary Table 2 for "Computational study and design of effective siRNAs to silence structural proteins associated genes of Indian SARS-CoV-2 strains"

| **Supplementary Table 2.** The conserved sequences of the Envelope gene identified from the 812 strains of SARS-CoV-2 | |
| --- | --- |
| **Position** | **Sequence** |
| 1-11 | ATGTACTCATT |
| 18-68 | GGAAGAGACAGGTACGTTAATAGTTAATAGCGTACTTCTTTTTCTTGCTTT |
| 70-95 | GTGGTATTCTTGCTAGTTACACTAGC |
| 97-109 | ATCCTTACTGCGC |
| 111-135 | TCGATTGTGTGCGTACTGCTGCAAT |
| 137-147 | TTGTTAACGTG |
| 149-156 | GTCTTGTA |
| 158-171 | AACCTTCTTTTTAC |
| 173-183 | TTTACTCTCGT |
| 185-224 | TTAAAAATCTGAATTCTTCTAGAGTTCCTGATCTTCTGGT |
