## Supplementary Table 3 for "Computational study and design of effective siRNAs to silence structural proteins associated genes of Indian SARS-CoV-2 strains"

| **Supplementary Table 3.** A conserved sequence of Membrane gene identified from the 812 strains of SARS-CoV-2. | |
| --- | --- |
| **Position** | **Sequences** |
| 1-4 | ATGG |
| 6-14 | AGATTCCAA |
| 16-83 | GGTACTATTACCGTTGAAGAGCTTAAAAAGCTCCTTGAACAATGGAACCTAGTAATAGGTTTCCTATT |
| 86-101 | TTACATGGATTTGTCT |
| 103-122 | CTACAATTTGCCTATGCCAA |
| 124-158 | AGGAATAGGTTTTTGTATATAATTAAGTTAATTTT |
| 160-167 | CTCTGGCT |
| 169-185 | TTATGGCCAGTAACTTT |
| 189-197 | TTGTTTTGT |
| 199-204 | CTTGCT |
| 214-244 | AGAATAAATTGGATCACCGGTGGAATTGCTA |
| 248-280 | CAATGGCTTGTCTTGTAGGCTTGATGTGGCTCA |
| 282-299 | CTACTTCATTGCTTCTTT |
| 321-388 | TTCCATGTGGTCATTCAA |
| 340-350 | CCAGAAACTAA |
| 352-359 | ATTCTTCT |
| 361-372 | AACGTGCCACTC |
| 374-384 | ATGGCACTATT |
| 386-401 | TGACCAGACCGCTTCT |
| 403-424 | GAAAGTGAACTCGTAATCGGAG |
| 430-482 | ATCCTTCGTGGACATCTTCGTATTGCTGGACACCATCTAGGACGCTGTGACAT |
| 484-491 | AAGGACCT |
| 493-605 | CCTAAAGAAATCACTGTTGCTACATCACGAACGCTTTCTTATTACAAATTGGGAGCTTCGCAGCGTGTAGCAGGTGACTCAGGTTTTGCTGCATACAGTCGCTACAGGATTGG |
| 610-624 | TATAAATTAAACACA |
| 626-641 | ACCATTCCAGTAGCAG |
| 643-669 | GACAATATTGCTTTGCTTGTACAGTAA |
