## Supplementary Table 4 for "Computational study and design of effective siRNAs to silence structural proteins associated genes of Indian SARS-CoV-2 strains"

| **Supplementary Table 4.** Conserved sequence of Nucleocapsid phosphoprotein gene identified from the 812 strains of SARS-CoV-2 | |
| --- | --- |
| **Position** | **Sequence** |
| 1-6 | ATGTCT |
| 10-15 | AATGGA |
| 17-26 | CCCAAAATCA |
| 28-33 | CGAAAT |
| 39-51 | CCGCATTACGTTT |
| 54-63 | TGGACCCTCA |
| 65-87 | ATTCAACTGGCAGTAACCAGAAT |
| 92-96 | AACGC |
| 109-116 | TCAAAACA |
| 118-122 | CGTCG |
| 124-142 | CCCCAAGGTTTACCCAATA |
| 144-164 | TACTGCGTCTTGGTTCACCGC |
| 166-198 | CTCACTCAACATGGCAAGGAAGACCTTAAATTC |
| 200-213 | CTCGAGGACAAGGC |
| 215-233 | TTCCAATTAACACCAATAG |
| 235-267 | AGTCCAGATGACCAAATTGGCTACTACCGAAGA |
| 269-275 | CTACCAG |
| 278-288 | GAATTCGTGGT |
| 290-308 | GTGACGGTAAAATGAAAGA |
| 310-329 | CTCAGTCCAAGATGGTATTT |
| 334-354 | TACCTAGGAACTGGGCCAGAA |
| 359-381 | GACTTCCCTATGGTGCTAACAAA |
| 383-400 | ACGGCATCATATGGGTTG |
| 405-414 | TGAGGGAGCC |
| 418-429 | AATACACCAAAA |
| 433-443 | CACATTGGCAC |
| 448-452 | CACATTGGCAC |
| 455-465 | CTAACAATGCT |
| 476-486 | TACAACTTCCT |
| 488-503 | AAGGAACAACATTGCC |
| 505-518 | AAAGGCTTCTACGC |
| 520-527 | GAAGGGAG |
| 529-537 | AGAGGCGGC |
| 540-571 | TCAAGCCTCTTCTCGTTCCTCATCACGTAGTC |
| 582-594 | AAGAAATTCAACT |
| 597-604 | AGGCAGCA |
| 615-625 | TTCTCCTGCTA |
| 631-652 | GCTGGCAATGGCGGTGATGCTG |
| 654-672 | TCTTGCTTTGCTGCTGCTT |
| 676-701 | AGATTGAACCAGCTTGAGAGCAAAAT |
| 707-711 | GTAAA |
| 717-749 | ACAACAACAAGGCCAAACTGTCACTAAGAAATC |
| 756-803 | TGAGGCTTCTAAGAAGCCTCGGCAAAAACGTACTGCCACTAAAGCATA |
| 809-860 | TAACACAAGCTTTCGGCAGACGTGGTCCAGAACAAACCCAAGGAAATTTTGG |
| 862-893 | GACCAGGAACTAATCAGACAAGGAACTGATTA |
| 899-903 | ATTGG |
| 907-914 | CAAATTGC |
| 916-926 | CAATTTGCCCC |
| 928-950 | AGCGCTTCAGCGTTCTTCGGAAT |
| 952-962 | TCGCGCATTGG |
| 964-977 | ATGGAAGTCACACC |
| 981-994 | GGGAACGTGGTTGA |
| 1021-1029 | GACAAAGAT |
| 1031-1041 | CAAATTTCAAA |
| 1043-1061 | ATCAAGTCATTTTGCTGAA |
| 1063-1074 | AAGCATATTGAC |
| 1086-1090 | ATTCC |
| 1092-1102 | ACCAACAGAGC |
| 1110-1121 | GGACAAAAAGAA |
| 1123-1128 | AAGGCT |
| 1130-1135 | ATGAAA |
| 1137-1153 | TCAAGCCTTACCGCAGA |
| 1155-1166 | ACAGAAGAAACA |
| 1170-1177 | AACTGTGA |
| 1179-1193 | TCTTCTTCCTGCTGC |
| 1221-1225 | GCAAC |
| 1227-1259 | ATCCATGAGCAGTGCTGACTCAACTCAGGCCTA |
