## Supplementary Table 5 for "Computational study and design of effective siRNAs to silence structural proteins associated genes of Indian SARS-CoV-2 strains"

| **Supplementary Table 5.** The conserved sequence of Surface Glycoprotein gene identified from the 812 strains of SARS-CoV-2. | |
| --- | --- |
| **Position** | **Sequence** |
| 39-51 | TCAGTGTGTTAAT |
| 55-64 | ACAACCAGAA |
| 69-79 | ATTACCCCCTG |
| 85-95 | ACTAATTCTTT |
| 97-103 | ACACGTG |
| 105-113 | TGTTTATTA |
| 171-175 | TTTCT |
| 181-193 | AATGTTACTTGGT |
| 195-202 | CCATGCTA |
| 217-223 | ACCAATG |
| 234-248 | GTTTGATAACCCTGT |
| 275-280 | TTGCTT |
| 290-327 | AGTCTAACATAATAAGAGGCTGGATTTTTGGTACTACT |
| 334-347 | TCGAAGACCCAGTC |
| 349-393 | CTACTTATTGTTAATAACGCTACTAATGTTGTTATTAAAGTCTGT |
| 397-410 | TTTCAATTTTGTAA |
| 413-418 | ATCCAT |
| 432-436 | TTACC |
| 443-449 | ACAACAA |
| 469-484 | TTCAGAGTTTATTCTA |
| 486-514 | TGCGAATAATTGCACTTTTGAATATGTCT |
| 520-530 | CCTTTTCTTAT |
| 532-541 | GACCTTGAAG |
| 543-555 | AAAACAGGGTAAT |
| 557-564 | TCAAAAAT |
| 566-617 | TTAGGGAATTTGTGTTTAAGAATATTGATGGTTATTTTAAAATATATTCTAA |
| 619-638 | CACACGCCTATTAATTTAGT |
| 640-656 | CGTGATCTCCCTCAGGG |
| 664-693 | GCTTTAGAACCATTGGTAGATTTGCCAATA |
| 698-710 | TTAACATCACTAG |
| 720-736 | TTTACTTGCTTTACATA |
| 740-746 | GTTATTT |
| 748-760 | ACTCCTGGTGATT |
| 774-780 | GACAGCT |
| 785-811 | CTGCAGCTTATTATGTGGGTTATCTTC |
| 815-821 | CTAGGAC |
| 823-869 | TTTCTATTAAAATATAATGAAAATGGAACCATTACAGATGCTGTAGA |
| 871-881 | TGTGCACTTGA |
| 889-901 | TCAGAAACAAAGT |
| 907-917 | TTGAAATCCTT |
| 919-1022 | GAATCTATTGTTAGATTTCCTAATATTACAAACTTGTGCCCTTTTGGTACTGTAGAAAAAGGAATCTATCAAACTTCTAACTTTAGAGTCCAACCAACAGAAGT |
| 1031-1036 | CCACCA |
| 1043-1055 | CATCTGTTTATGC |
| 1057-1062 | TGGAAC |
| 1064-1082 | GGAAGAGAATCAGCAACTG |
| 1084-1115 | GTTGCTGATTATTCTGTCCTATATAATTCCGC |
| 1117-1138 | TCATTTTCCACTTTTAAGTGTT |
| 1216-1222 | GAAGTCA |
| 1224-1298 | ACAAATCGCTCCAGGGCAAACTGGAAAGATTGCTGATTATAATTATAAATTACCAGATGATTTTACAGGCTGCGT |
| 1300-1330 | ATAGCTTGGAATTCTAACAATCTTGATTCTA |
| 1332-1354 | GGTTGGTGGTAATTATAATTACC |
| 1356-1373 | GTATAGATTGTTTAGGAA |
| 1375-1401 | TCTAATCTCAAACCTTTTGAGAGAGAT |
| 1403-1410 | TTTCAACT |
| 1415-1424 | TCTATCAGGC |
| 1435-1449 | CCTTGTAATGGTGTT |
| 1453-1468 | GGTTTTAATTGTTACT |
| 1470-1500 | TCCTTTACAATCATATGGTTTCCAACCCACT |
| 1502-1517 | ATGGTGTTGGTTACCA |
| 1523-1553 | ACAGAGTAGTAGTACTTTCTTTTGAACTTCT |
| 1555-1601 | CATGCACCAGCAACTGTTTGTGGACCTAAAAAGTCTACTAATTTGGT |
| 1603-1667 | AAAAACAAATGTGTCAATTTCAACTTCAATGGTTTAACAGGCACAGGTGTTCTTACTGAGTCTAA |
| 1675-1707 | TTTCTGCCTTTCCAACAATTTGGCAGAGACATT |
| 1710-1714 | TGACA |
| 1721-1748 | ATGCTGTCCGTGATCCACAGACACTTGA |
| 1750-1838 | ATTCTTGACATTACACCATGTTCTTTTGGTGGTGTCAGTGTTATAACACCAGGAACAAATACTTCTAACCAGGTTGCTGTTCTTTATCA |
| 1842-1863 | TGTTAACTGCACAGAAGTCCCT |
| 1865-1878 | TTGCTATTCATGCA |
| 1881-1925 | TCAACTTACTCCTACTTGGCGTGTTTATTCTACAGGTTCTAATGT |
| 1927-1943 | TTTCAAACACGTGCAGG |
| 1945-1962 | TGTTTAATAGGGGCTGAA |
| 1964-2024 | ATGTCAACAACTCATATGAGTGTGACATACCCATTGGTGCAGGTATATGCGCTAGTTATCA |
| 2026-2030 | ACTCA |
| 2032-2041 | ACTAATTCTC |
| 2050-2061 | GCACGTAGTGTA |
| 2077-2100 | ATTGCCTACACTATGTCACTTGGT |
| 2103-2110 | AGAAAATT |
| 2117-2130 | CTTACTCTAATAAC |
| 2132-2146 | CTATTGCCATACCCA |
| 2148-2192 | AAATTTTACTATTAGTGTTACCACAGAAATTCTACCAGTGTCTAT |
| 2194-2339 | ACCAAGACATCAGTAGATTGTACAATGTACATTTGTGGTGATTCAACTGAATGCAGCAATCTTTTGTTGCAATATGGCAGTTTTTGTACACAATTAAACCGTGCTTTAACTGGAATAGCTGTTGAACAAGACAAAAACACCCAAGA |
| 2341-2348 | GTTTTTGC |
| 2368-2382 | AAAACACCACCAATT |
| 2386-2412 | GATTTTGGTGGTTTTAATTTTTCACAA |
| 2450-2464 | TTATTGAAGATCTAC |
| 2466-2479 | TTTCAACAAAGTGA |
| 2482-2519 | CTTGCAGATGCTGGCTTCATCAAACAATATGGTGATTG |
| 2521-2532 | CTTGGTGATATT |
| 2534-2567 | CTGCTAGAGACCTCATTTGTGCACAAAAGTTTAA |
| 2569-2611 | GGCCTTACTGTTTTGCCACCTTTGCTCACAGATGAAATGATTG |
| 2613-2628 | TCAATACACTTCTGCA |
| 2636-2642 | CGGGTAC |
| 2644-2674 | ATCACTTCTGGTTGGACCTTTGGTGCAGGTG |
| 2676-2720 | TGCATTACAAATACCATTTGCTATGCAAATGGCTTATAGGTTTAA |
| 2722-2785 | GGTATTGGAGTTACACAGAATGTTCTCTATGAGAACCAAAAATTGATTGCCAACCAATTTAATA |
| 2790-2795 | TATTGG |
| 2797-2805 | AAAATTCAA |
| 2809-2815 | TCACTTT |
| 2817-2826 | TTCCACAGCA |
| 2829-2903 | TGCACTTGGAAAACTTCAAGATGTGGTCAACCAAAATGCACAAGCTTTAAACACGCTTGTTAAACAACTTAGCTC |
| 2905-2939 | AATTTTGGTGCAATTTCAAGTGTTTTAAATGATAT |
| 2945-2972 | CACGTCTTGACAAAGTTGAGGCTGAAGT |
| 2974-2987 | CAAATTGATAGGTT |
| 2992-3110 | ACAGGCAGACTTCAAAGTTTGCAGACATATGTGACTCAACAATTAATTAGAGCTGCAGAAATCAGAGCTTCTGCTAATCTTGCTGCTACTAAAATGTCAGAGTGTGTACTTGGACAATC |
| 3112-3134 | AAAAGAGTTGATTTTTGTGGAAA |
| 3136-3149 | GGCTATCATCTTAT |
| 3151-3158 | TCCTTCCC |
| 3160-3179 | CAGTCAGCACCTCATGGTGT |
| 3181-3188 | GTCTTCTT |
| 3190-3194 | CATGT |
| 3198-3211 | TTATGTCCCTGCAC |
| 3214-3221 | GAAAAGAA |
| 3223-3229 | TTCACAA |
| 3232-3248 | GCTCCTGCCATTTGTCA |
| 3251-3268 | ATGGAAAAGCAC |
| 3266-3271 | TTCCTC |
| 3273-3302 | TGAAGGTGTCTTTGTTTCAAATGGCACACA |
| 3304-3309 | TGGTTT |
| 3311-3329 | TAACACAAAGGAATTTTTA |
| 3331-3345 | GAACCACAAATCATT |
| 3347-3351 | CTACA |
| 3358-3363 | ACATTT |
| 3365-3383 | TGTCTGGTAACTGTGATGT |
| 3385-3427 | GTAATAGGAATTGTCAACAACACAGTTTATGATCCTTTGCAAC |
| 3429-3443 | TGAATTAGACTCATT |
| 3445-3456 | AAGGAGGAGTTA |
| 3458-3476 | ATAAATATTTTAAGAATCA |
| 3478-3486 | ACATCACCA |
| 3488-3525 | ATGTTGATTTAGGTGACATCTCTGGCATTAATGCTTCA |
| 3527-3538 | TTGTAAACATTC |
| 3548-3560 | TTGACCGCCTCAA |
| 3562-3572 | GAGGTTGCCAA |
| 3576-3599 | TTTAAATGAATCTCTCATCGATCT |
| 3602-3647 | AAGAACTTGGAAAGTATGAGCAGTATATAAAATGGCCATGGTACAT |
| 3649-3713 | TGGCTAGGTTTTATAGCTGGCTTGATTGCCATAGTAATGGTGACAATTATGCTTTGCTGTATGAC |
| 3715-3727 | AGTTGCTGTAGTT |
| 3729-3761 | TCTCAAGGGCTGTTGTTCTTGTGGATCCTGCTG |
| 3763-3774 | AAATTTGATGAA |
| 3776-3787 | ACGACTCTGAGC |
| 3793-3821 | CTCAAAGGAGTCAAATTACATTACACATA |
