## Supplementary Table 6 for "Computational study and design of effective siRNAs to silence structural proteins associated genes of Indian SARS-CoV-2 strains"

**Supplementary Table 6.** Tm values of predicted siRNA (guide strand and passenger strand) against envelope gene.

| **S.no** | **Alias** | **Conserved position** | **Location of target within mRNA** | **siRNA target within mRNA** | **Predicted siRNA siRNA duplex candidate at 37◦C; RNA oligo sequences  21nt guide (5' →3’)**  **21nt passenger (5' →3’)** | **Functional siRNA selection: Ui-Tei, Reynolds and Amarzguioui** | **Seed duplex (Tm) guide** | **Seed duplex (Tm) Passenger** |
| --- | --- | --- | --- | --- | --- | --- | --- | --- |
| 1 | e1 | 18-68 | 13-35 | TACGTTAATAGTTAATAGCGTAC | ACGCUAUUAACUAUUAACGUA CGUUAAUAGUUAAUAGCGUAC | **U A** | 20.4 °C | 1.4 °C |
| 2 | e2 | 18-68 | 21-43 | TAGTTAATAGCGTACTTCTTTTT | AAAGAAGUACGCUAUUAACUA GUUAAUAGCGUACUUCUUUUU | **U A** | 17.7 °C | -2.3 °C |
| 3 | e3 | 18-68 | 22-44 | AGTTAATAGCGTACTTCTTTTTC | AAAAGAAGUACGCUAUUAACU UUAAUAGCGUACUUCUUUUUC | **R** | 10.3 °C | 14.9 °C |
| 4 | e4 | 18-68 | 23-45 | GTTAATAGCGTACTTCTTTTTCT | AAAAAGAAGUACGCUAUUAAC UAAUAGCGUACUUCUUUUUCU | **R** | 5.5 °C | 20.4 °C |
| 5 | e5 | 70-95 | 1-23 | GTGGTATTCTTGCTAGTTACACT | UGUAACUAGCAAGAAUACCAC GGUAUUCUUGCUAGUUACACU | **U R A** | 14.3 °C | 14.5 °C |
| 6 | e6 | 185-224 | 1-23 | TTAAAAATCTGAATTCTTCTAGA | UAGAAGAAUUCAGAUUUUUAA AAAAAUCUGAAUUCUUCUAGA | **R** | 19.1 °C | 5.3 °C |
| 7 | e7 | 185-224 | 3-25 | AAAAATCTGAATTCTTCTAGAGT | UCUAGAAGAAUUCAGAUUUUU AAAUCUGAAUUCUUCUAGAGU | **R** | 16.3 °C | 20.4 °C |
