## Supplementary Table 7 for "Computational study and design of effective siRNAs to silence structural proteins associated genes of Indian SARS-CoV-2 strains"

**Supplementary Table 7.** Tm values of predicted siRNA (guide strand and passenger strand) against Membrane gene.

| **S.no** | **Alias** | **Conserved position** | **Location of target within mRNA** | **siRNA target within mRNA** | **Predicted siRNA siRNA duplex candidate at 37◦C; RNA oligo sequences  21nt guide (5' →3')**  **21nt passenger (5' →3')** | **Functional siRNA selection: Ui-Tei, Reynolds and Amarzguioui** | **Seed duplex (Tm) guide** | **Seed duplex (Tm) Passenger** |
| --- | --- | --- | --- | --- | --- | --- | --- | --- |
| 1 | m1 | 16-83 | 9-31 | TACCGTTGAAGAGCTTAAAAAGC | UUUUUAAGCUCUUCAACGGUA CCGUUGAAGAGCUUAAAAAGC | U R A | -3.8 °C | 21.1 °C |
| 2 | m2 | 16-83 | 21-43 | GCTTAAAAAGCTCCTTGAACAAT | UGUUCAAGGAGCUUUUUAAGC UUAAAAAGCUCCUUGAACAAU | R | 19.2 °C | -3.8 °C |
| 3 | m3 | 16-83 | 22-44 | CTTAAAAAGCTCCTTGAACAATG | UUGUUCAAGGAGCUUUUUAAG UAAAAAGCUCCUUGAACAAUG | R | 20.5 °C | 12.3 °C |
| 4 | m4 | 124-158 | 1-23 | AGGAATAGGTTTTTGTATATAAT | UAUAUACAAAAACCUAUUCCU GAAUAGGUUUUUGUAUAUAAU | U R A | 8.2 °C | 18.5 °C |
| 5 | m5 | 124-158 | 2-24 | GGAATAGGTTTTTGTATATAATT | UUAUAUACAAAAACCUAUUCC AAUAGGUUUUUGUAUAUAAUU | R | 2.8 °C | 18.5 °C |
| 6 | m6 | 124-158 | 3-25 | GAATAGGTTTTTGTATATAATTA | AUUAUAUACAAAAACCUAUUC AUAGGUUUUUGUAUAUAAUUA | R | -5.9 °C | 18.6 °C |
| 7 | m7 | 124-158 | 5-27 | ATAGGTTTTTGTATATAATTAAG | UAAUUAUAUACAAAAACCUAU AGGUUUUUGUAUAUAAUUAAG | R A | -8.0 °C | 12.6 °C |
| 8 | m8 | 124-158 | 6-28 | TAGGTTTTTGTATATAATTAAGT | UUAAUUAUAUACAAAAACCUA GGUUUUUGUAUAUAAUUAAGU | U R A | -8.0 °C | 5.6 °C |
| 9 | m9 | 124-158 | 12-34 | TTTGTATATAATTAAGTTAATTT | AUUAACUUAAUUAUAUACAAA UGUAUAUAAUUAAGUUAAUUU | R | 4.9 °C | 2.8 °C |
| 10 | m10 | 124-158 | 13-35 | TTGTATATAATTAAGTTAATTTT | AAUUAACUUAAUUAUAUACAA GUAUAUAAUUAAGUUAAUUUU | U R A | 4.6 °C | -5.9 °C |
