## Supplementary Table 8 for "Computational study and design of effective siRNAs to silence structural proteins associated genes of Indian SARS-CoV-2 strains"

**Supplementary Table 8.** Tm values of predicted siRNA (guide strand and passenger strand) against nucleocapsid phosphoprotein gene.

| **S.no** | **Alias** | **Conserved position** | **Location of target within mRNA** | **siRNA target within mRNA** | **Predicted siRNA siRNA duplex candidate at 37◦C; RNA oligo sequences  21nt guide (5' →3’)**  **21nt passenger (5' →3’)** | **Functional siRNA selection: Ui-Tei, Reynolds and Amarzguioui** | **Seed duplex (Tm) guide** | **Seed duplex (Tm) Passenger** |
| --- | --- | --- | --- | --- | --- | --- | --- | --- |
| 1 | n1 | 717-749 | 11-33 | GGCCAAACTGTCACTAAGAAATC | UUUCUUAGUGACAGUUUGGCC CCAAACUGUCACUAAGAAAUC | **U R A** | 11.7 °C | 16.7 °C |
| 2 | n2 | 809-860 | 27-49 | CCAGAACAAACCCAAGGAAATTT | AUUUCCUUGGGUUUGUUCUGG AGAACAAACCCAAGGAAAUUU | **R A** | 18.7 °C | 14.9 °C |
| 3 | n3 | 809-860 | 28-50 | CAGAACAAACCCAAGGAAATTTT | AAUUUCCUUGGGUUUGUUCUG GAACAAACCCAAGGAAAUUUU | **U R A** | 18.7 °C | 13.3 °C |
