## Supplementary Table 9 for "Computational study and design of effective siRNAs to silence structural proteins associated genes of Indian SARS-CoV-2 strains"

**Supplementary Table 9.** Tm values of predicted siRNA (guide strand and passenger strand) against surface glycoprotein gene.

| **S.no** | **Alias** | **Conserved position** | **Location of target within mRNA** | **siRNA target within mRNA** | **Predicted siRNA siRNA duplex candidate at 37◦C; RNA oligo sequences  21nt guide (5' →3' ) 21nt passenger (5' →3' )** | **Functional siRNA selection: Ui-Tei, Reynolds and Amarzguioui** | **Seed duplex (Tm) guide** | **Seed duplex (Tm) Passenger** |
| --- | --- | --- | --- | --- | --- | --- | --- | --- |
| 1 | g1 | 349-393 | 10-32 | GTTAATAACGCTACTAATGTTGT | AACAUUAGUAGCGUUAUUAAC UAAUAACGCUACUAAUGUUGU | **R** | 11.6 °C | 6.9 °C |
| 2 | g2 | 349-393 | 17-39 | ACGCTACTAATGTTGTTATTAAA | UAAUAACAACAUUAGUAGCGU GCUACUAAUGUUGUUAUUAAA | **UA** | 6.9 °C | 11.3 °C |
| 3 | g3 | 349-393 | 18-40 | CGCTACTAATGTTGTTATTAAAG | UUAAUAACAACAUUAGUAGCG CUACUAAUGUUGUUAUUAAAG | **URA** | 1.4 °C | 6.3 °C |
| 4 | g4 | 349-393 | 19-41 | GCTACTAATGTTGTTATTAAAGT | UUUAAUAACAACAUUAGUAGC UACUAAUGUUGUUAUUAAAGU | **R** | -7.5 °C | 11.6 °C |
| 5 | g5 | 349-393 | 21-43 | TACTAATGTTGTTATTAAAGTCT | ACUUUAAUAACAACAUUAGUA CUAAUGUUGUUAUUAAAGUCU | **UR** | -4.3 °C | 6.9 °C |
| 6 | g6 | 486-514 | 1-23 | TGCGAATAATTGCACTTTTGAAT | UCAAAAGUGCAAUUAUUCGCA CGAAUAAUUGCACUUUUGAAU | **URA** | 10.3 °C | 1.8 °C |
| 7 | g7 | 486-514 | 2-24 | GCGAATAATTGCACTTTTGAATA | UUCAAAAGUGCAAUUAUUCGC GAAUAAUUGCACUUUUGAAUA | **URA** | 12.2 °C | -10.3 °C |
| 8 | g8 | 486-514 | 4-26 | GAATAATTGCACTTTTGAATATG | UAUUCAAAAGUGCAAUUAUUC  AUAAUUGCACUUUUGAAUAUG | **R** | 7.4 °C | 15.3 °C |
| 9 | g9 | 486-514 | 5-27 | AATAATTGCACTTTTGAATATGT | AUAUUCAAAAGUGCAAUUAUU  UAAUUGCACUUUUGAAUAUGU | **R** | 8.9 °C | 20.0 °C |
| 10 | g10 | 566-617 | 2-24 | TAGGGAATTTGTGTTTAAGAATA | UUCUUAAACACAAAUUCCCUA GGGAAUUUGUGUUUAAGAAUA | **URA** | 7.1 °C | 14.2 °C |
| 11 | g11 | 566-617 | 3-25 | AGGGAATTTGTGTTTAAGAATAT | AUUCUUAAACACAAAUUCCCU  GGAAUUUGUGUUUAAGAAUAU | **UA** | 7.1 °C | 7.4 °C |
| 12 | g12 | 566-617 | 4-26 | GGGAATTTGTGTTTAAGAATATT | UAUUCUUAAACACAAAUUCCC  GAAUUUGUGUUUAAGAAUAUU | **URA** | 6.9 °C | 5.3 °C |
| 13 | g13 | 566-617 | 5-27 | GGAATTTGTGTTTAAGAATATTG | AUAUUCUUAAACACAAAUUCC  AAUUUGUGUUUAAGAAUAUUG | **R** | 6.9 °C | 12.1 °C |
| 14 | g14 | 566-617 | 15-37 | TTTAAGAATATTGATGGTTATTT | AUAACCAUCAAUAUUCUUAAA  UAAGAAUAUUGAUGGUUAUUU | **R** | 20.0 °C | 6.9 °C |
| 15 | g15 | 566-617 | 17-39 | TAAGAATATTGATGGTTATTTTA | AAAUAACCAUCAAUAUUCUUA  AGAAUAUUGAUGGUUAUUUUA | **A** | 13.9 °C | 1.8 °C |
| 16 | g16 | 566-617 | 18-40 | AAGAATATTGATGGTTATTTTAA | AAAAUAACCAUCAAUAUUCUU  GAAUAUUGAUGGUUAUUUUAA | **URA** | -0.3 °C | -1.8 °C |
| 17 | g17 | 566-617 | 19-41 | AGAATATTGATGGTTATTTTAAA | UAAAAUAACCAUCAAUAUUCU  AAUAUUGAUGGUUAUUUUAAA | **R** | -9.7 °C | 8.7 °C |
| 18 | g18 | 566-617 | 20-42 | GAATATTGATGGTTATTTTAAAA | UUAAAAUAACCAUCAAUAUUC  AUAUUGAUGGUUAUUUUAAAA | **R** | -7.5 °C | 8.7 °C |
| 19 | g19 | 566-617 | 21-43 | AATATTGATGGTTATTTTAAAAT | UUUAAAAUAACCAUCAAUAUU  UAUUGAUGGUUAUUUUAAAAU | **R** | -9.7 °C | 13.6 °C |
| 20 | g20 | 566-617 | 25-47 | TTGATGGTTATTTTAAAATATAT | AUAUUUUAAAAUAACCAUCAA  GAUGGUUAUUUUAAAAUAUAU | **URA** | -7.5 °C | 20.0 °C |
| 21 | g21 | 566-617 | 26-48 | TGATGGTTATTTTAAAATATATT | UAUAUUUUAAAAUAACCAUCA  AUGGUUAUUUUAAAAUAUAUU | **R** | -10.3 °C | 20.0 °C |
| 22 | g22 | 566-617 | 27-49 | GATGGTTATTTTAAAATATATTC | AUAUAUUUUAAAAUAACCAUC UGGUUAUUUUAAAAUAUAUUC | **RA** | -8.0 °C | 13.9 °C |
| 23 | g23 | 566-617 | 28-50 | ATGGTTATTTTAAAATATATTCT | AAUAUAUUUUAAAAUAACCAU GGUUAUUUUAAAAUAUAUUCU | **UA** | -8.6 °C | -0.3 °C |
| 24 | g24 | 823-869 | 2-24 | TTCTATTAAAATATAATGAAAAT | UUUCAUUAUAUUUUAAUAGAA CUAUUAAAAUAUAAUGAAAAU | **URA** | 8.9 °C | -7.5 °C |
| 25 | g25 | 823-869 | 4-26 | CTATTAAAATATAATGAAAATGG | AUUUUCAUUAUAUUUUAAUAG  AUUAAAAUAUAAUGAAAAUGG | **R** | 7.4 °C | -9.7 °C |
| 26 | g26 | 823-869 | 7-29 | TTAAAATATAATGAAAATGGAAC | UCCAUUUUCAUUAUAUUUUAA  AAAAUAUAAUGAAAAUGGAAC | **R** | 11.3 °C | -8.0 °C |
| 27 | g27 | 823-869 | 16-38 | AATGAAAATGGAACCATTACAGA | UGUAAUGGUUCCAUUUUCAUU  UGAAAAUGGAACCAUUACAGA | **R** | 20.0 °C | 7.4 °C |
| 28 | g28 | 919-1022 | 3-25 | ATCTATTGTTAGATTTCCTAATA | UUAGGAAAUCUAACAAUAGAU  CUAUUGUUAGAUUUCCUAAUA | **UA** | 19.9 °C | 6.9 °C |
| 29 | g29 | 919-1022 | 5-27 | CTATTGTTAGATTTCCTAATATT | UAUUAGGAAAUCUAACAAUAG  AUUGUUAGAUUUCCUAAUAUU | **R** | 19.9 °C | 11.8 °C |
| 30 | g30 | 919-1022 | 6-28 | TATTGTTAGATTTCCTAATATTA | AUAUUAGGAAAUCUAACAAUA  UUGUUAGAUUUCCUAAUAUUA | **R** | 12.3 °C | 20.3 °C |
| 31 | g31 | 919-1022 | 7-29 | ATTGTTAGATTTCCTAATATTAC | AAUAUUAGGAAAUCUAACAAU  UGUUAGAUUUCCUAAUAUUAC | **R** | -2.7 °C | 14.5 °C |
| 32 | g32 | 919-1022 | 8-30 | TTGTTAGATTTCCTAATATTACA | UAAUAUUAGGAAAUCUAACAA  GUUAGAUUUCCUAAUAUUACA | **URA** | -8.0 °C | 6.9 °C |
| 33 | g33 | 919-1022 | 10-32 | GTTAGATTTCCTAATATTACAAA | UGUAAUAUUAGGAAAUCUAAC  UAGAUUUCCUAAUAUUACAAA | **R** | 1.1 °C | 14.8 °C |
| 34 | g34 | 919-1022 | 12-34 | TAGATTTCCTAATATTACAAACT | UUUGUAAUAUUAGGAAAUCUA  GAUUUCCUAAUAUUACAAACU | **UA** | 6.9 °C | 18.7 °C |
| 35 | g35 | 919-1022 | 17-39 | TTCCTAATATTACAAACTTGTGC | ACAAGUUUGUAAUAUUAGGAA  CCUAAUAUUACAAACUUGUGC | **UA** | 10.3 °C | -2.7 °C |
| 36 | g36 | 919-1022 | 38-60 | GCCCTTTTGGTACTGTAGAAAAA | UUUCUACAGUACCAAAAGGGC  CCUUUUGGUACUGUAGAAAAA | **UA** | 20.3 °C | 16.1 °C |
| 37 | g37 | 919-1022 | 39-61 | CCCTTTTGGTACTGTAGAAAAAG | UUUUCUACAGUACCAAAAGGG  CUUUUGGUACUGUAGAAAAAG | **UA** | 14.6 °C | 18.8 °C |
| 38 | g38 | 919-1022 | 40-62 | CCTTTTGGTACTGTAGAAAAAGG | UUUUUCUACAGUACCAAAAGG  UUUUGGUACUGUAGAAAAAGG | **R** | 7.1 °C | 20.0 °C |
| 39 | g39 | 919-1022 | 45-67 | TGGTACTGTAGAAAAAGGAATCT | AUUCCUUUUUCUACAGUACCA  GUACUGUAGAAAAAGGAAUCU | **UA** | 18.7 °C | 20.2 °C |
| 40 | g40 | 919-1022 | 52-74 | GTAGAAAAAGGAATCTATCAAAC | UUGAUAGAUUCCUUUUUCUAC  AGAAAAAGGAAUCUAUCAAAC | **R** | 21.4 °C | 5.5 °C |
| 41 | g41 | 919-1022 | 53-75 | TAGAAAAAGGAATCTATCAAACT | UUUGAUAGAUUCCUUUUUCUA  GAAAAAGGAAUCUAUCAAACU | **UA** | 13.4 °C | 9.5 °C |
| 42 | g42 | 919-1022 | 56-78 | AAAAAGGAATCTATCAAACTTCT | AAGUUUGAUAGAUUCCUUUUU  AAAGGAAUCUAUCAAACUUCU | **R** | 19.2 °C | 18.7 °C |
| 43 | g43 | 919-1022 | 59-81 | AAGGAATCTATCAAACTTCTAAC | UAGAAGUUUGAUAGAUUCCUU  GGAAUCUAUCAAACUUCUAAC | **URA** | 17.7 °C | 16.0 °C |
| 44 | g44 | 919-1022 | 60-82 | AGGAATCTATCAAACTTCTAACT | UUAGAAGUUUGAUAGAUUCCU  GAAUCUAUCAAACUUCUAACU | **URA** | 18.9 °C | 6.6 °C |
| 45 | g45 | 919-1022 | 64-86 | ATCTATCAAACTTCTAACTTTAG | AAAGUUAGAAGUUUGAUAGAU  CUAUCAAACUUCUAACUUUAG | **URA** | 9.8 °C | 8.9 °C |
| 46 | g46 | 919-1022 | 65-87 | TCTATCAAACTTCTAACTTTAGA | UAAAGUUAGAAGUUUGAUAGA  UAUCAAACUUCUAACUUUAGA | **R** | 4.9 °C | 14.8 °C |
| 47 | g47 | 919-1022 | 67-89 | TATCAAACTTCTAACTTTAGAGT | UCUAAAGUUAGAAGUUUGAUA  UCAAACUUCUAACUUUAGAGU | **R** | 9.8 °C | 10.3 °C |
| 48 | g48 | 1084-1115 | 4-26 | GCTGATTATTCTGTCCTATATAA | AUAUAGGACAGAAUAAUCAGC  UGAUUAUUCUGUCCUAUAUAA | **R** | 21.0 °C | 1.8 °C |
| 49 | g49 | 1084-1115 | 5-27 | CTGATTATTCTGTCCTATATAAT | UAUAUAGGACAGAAUAAUCAG  GAUUAUUCUGUCCUAUAUAAU | **URA** | 12.2 °C | 1.8 °C |
| 50 | g50 | 1084-1115 | 6-28 | TGATTATTCTGTCCTATATAATT | UUAUAUAGGACAGAAUAAUCA  AUUAUUCUGUCCUAUAUAAUU | **R** | -0.8 °C | 6.9 °C |
| 51 | g51 | 1084-1115 | 7-29 | GATTATTCTGTCCTATATAATTC | AUUAUAUAGGACAGAAUAAUC  UUAUUCUGUCCUAUAUAAUUC | **R** | -5.9 °C | 13.4 °C |
| 52 | g52 | 1224-1298 | 21-43 | CTGGAAAGATTGCTGATTATAAT | UAUAAUCAGCAAUCUUUCCAG  GGAAAGAUUGCUGAUUAUAAU | **URA** | 8.7 °C | 14.8 °C |
| 53 | g53 | 1224-1298 | 22-44 | TGGAAAGATTGCTGATTATAATT | UUAUAAUCAGCAAUCUUUCCA  GAAAGAUUGCUGAUUAUAAUU | **URA** | 3.5 °C | 5.3 °C |
| 54 | g54 | 1224-1298 | 23-45 | GGAAAGATTGCTGATTATAATTA | AUUAUAAUCAGCAAUCUUUCC  AAAGAUUGCUGAUUAUAAUUA | **R** | -8.0 °C | 12.0 °C |
| 55 | g55 | 1224-1298 | 30-52 | TTGCTGATTATAATTATAAATTA | AUUUAUAAUUAUAAUCAGCAA GCUGAUUAUAAUUAUAAAUUA | **UA** | -7.5 °C | 13.4 °C |
| 56 | g56 | 1224-1298 | 31-53 | TGCTGATTATAATTATAAATTAC | AAUUUAUAAUUAUAAUCAGCA  CUGAUUAUAAUUAUAAAUUAC | **UA** | -8.0 °C | 8.7 °C |
| 57 | g57 | 1224-1298 | 32-54 | GCTGATTATAATTATAAATTACC | UAAUUUAUAAUUAUAAUCAGC  UGAUUAUAAUUAUAAAUUACC | **R** | -10.3 °C | 3.5 °C |
| 58 | g58 | 1224-1298 | 35-57 | GATTATAATTATAAATTACCAGA | UGGUAAUUUAUAAUUAUAAUC  UUAUAAUUAUAAAUUACCAGA | **R** | 13.9 °C | -8.0 °C |
| 59 | g59 | 1224-1298 | 44-66 | TATAAATTACCAGATGATTTTAC | AAAAUCAUCUGGUAAUUUAUA UAAAUUACCAGAUGAUUUUAC | **R** | 7.2 °C | -0.3 °C |
| 60 | g60 | 1224-1298 | 45-67 | ATAAATTACCAGATGATTTTACA | UAAAAUCAUCUGGUAAUUUAU AAAUUACCAGAUGAUUUUACA | **R** | 7.4 °C | 13.9 °C |
| 61 | g61 | 1300-1330 | 3-25 | AGCTTGGAATTCTAACAATCTTG | AGAUUGUUAGAAUUCCAAGCU  CUUGGAAUUCUAACAAUCUUG | **UA** | 14.8 °C | 20.1 °C |
| 62 | g62 | 1300-1330 | 6-28 | TTGGAATTCTAACAATCTTGATT | UCAAGAUUGUUAGAAUUCCAA  GGAAUUCUAACAAUCUUGAUU | **UA** | 12.0 °C | 14.8 °C |
| 63 | g63 | 1300-1330 | 7-29 | TGGAATTCTAACAATCTTGATTC | AUCAAGAUUGUUAGAAUUCCA  GAAUUCUAACAAUCUUGAUUC | **U** | 20.4 °C | 6.9 °C |
| 64 | g64 | 1470-1500 | 1-23 | TCCTTTACAATCATATGGTTTCC | AAACCAUAUGAUUGUAAAGGA CUUUACAAUCAUAUGGUUUCC | **UA** | 20.0 °C | 7.2 °C |
| 65 | g65 | 1523-1553 | 2-24 | CAGAGTAGTAGTACTTTCTTTTG | AAAGAAAGUACUACUACUCUG  GAGUAGUAGUACUUUCUUUUG | **UA** | 10.3 °C | 18.8 °C |
| 66 | g66 | 1523-1553 | 5-27 | AGTAGTAGTACTTTCTTTTGAAC | UCAAAAGAAAGUACUACUACU UAGUAGUACUUUCUUUUGAAC | **R** | 12.2 °C | 18.8 °C |
| 67 | g67 | 1523-1553 | 6-28 | GTAGTAGTACTTTCTTTTGAACT | UUCAAAAGAAAGUACUACUAC AGUAGUACUUUCUUUUGAACU | **R** | 12.2 °C | 21.3 °C |
| 68 | g68 | 1555-1601 | 11-33 | CAACTGTTTGTGGACCTAAAAAG | UUUUAGGUCCACAAACAGUUG ACUGUUUGUGGACCUAAAAAG | **R** | 18.6 °C | 16.7 °C |
| 69 | g69 | 1555-1601 | 12-34 | AACTGTTTGTGGACCTAAAAAGT | UUUUUAGGUCCACAAACAGUU CUGUUUGUGGACCUAAAAAGU | **URA** | 11.0 °C | 19.3 °C |
| 70 | g70 | 1555-1601 | 21-43 | TGGACCTAAAAAGTCTACTAATT | UUAGUAGACUUUUUAGGUCCA GACCUAAAAAGUCUACUAAUU | **UA** | 20.1 °C | 18.6 °C |
| 71 | g71 | 1555-1601 | 22-44 | GGACCTAAAAAGTCTACTAATTT | AUUAGUAGACUUUUUAGGUCC ACCUAAAAAGUCUACUAAUUU | **A** | 11.3 °C | 11.0 °C |
| 72 | g72 | 1555-1601 | 23-45 | GACCTAAAAAGTCTACTAATTTG | AAUUAGUAGACUUUUUAGGUC CCUAAAAAGUCUACUAAUUUG | **URA** | 6.3 °C | -3.8 °C |
| 73 | g73 | 1555-1601 | 24-46 | ACCTAAAAAGTCTACTAATTTGG | AAAUUAGUAGACUUUUUAGGU CUAAAAAGUCUACUAAUUUGG | **URA** | 4.6 °C | -3.8 °C |
| 74 | g74 | 1603-1667 | 3-25 | AAACAAATGTGTCAATTTCAACT | UUGAAAUUGACACAUUUGUUU ACAAAUGUGUCAAUUUCAACU | **R** | 7.4 °C | 12.1 °C |
| 75 | g75 | 1603-1667 | 9-31 | ATGTGTCAATTTCAACTTCAATG | UGAAGUUGAAAUUGACACAU GUGUCAAUUUCAACUUCAAUG | **URA** | 19.2 °C | 20.5 °C |
| 76 | g76 | 1603-1667 | 19-41 | TTCAACTTCAATGGTTTAACAGG | UGUUAAACCAUUGAAGUUGAA CAACUUCAAUGGUUUAACAGG | **UR** | 8.2 °C | 19.2 °C |
| 77 | g77 | 1750-1838 | 5-27 | TTGACATTACACCATGTTCTTTT | AAGAACAUGGUGUAAUGUCAA GACAUUACACCAUGUUCUUUU | **UA** | 19.2 °C | 14.6 °C |
| 78 | g78 | 1750-1838 | 6-28 | TGACATTACACCATGTTCTTTTG | AAAGAACAUGGUGUAAUGUCA ACAUUACACCAUGUUCUUUUG | **A** | 19.2 °C | 13.5 °C |
| 79 | g79 | 1750-1838 | 7-29 | GACATTACACCATGTTCTTTTGG | AAAAGAACAUGGUGUAAUGUC CAUUACACCAUGUUCUUUUGG | **UA** | 13.3 °C | 14.6 °C |
| 80 | g80 | 1881-1925 | 9-31 | CTCCTACTTGGCGTGTTTATTCT | AAUAAACACGCCAAGUAGGAG CCUACUUGGCGUGUUUAUUCU | **UA** | 6.9 °C | 16.4 °C |
| 81 | g81 | 1964-2024 | 1-23 | ATGTCAACAACTCATATGAGTGT | ACUCAUAUGAGUUGUUGACAU GUCAACAACUCAUAUGAGUGU | **UA** | 13.3 °C | 20.5 °C |
| 82 | g82 | 1964-2024 | 39-61 | GCAGGTATATGCGCTAGTTATCA | AUAACUAGCGCAUAUACCUGC AGGUAUAUGCGCUAGUUAUCA | **A** | 11.3 °C | 15.2 °C |
| 83 | g83 | 2194-2339 | 1-23 | ACCAAGACATCAGTAGATTGTAC | ACAAUCUACUGAUGUCUUGGU CAAGACAUCAGUAGAUUGUAC | **UR** | 13.4 °C | 19.2 °C |
| 84 | g84 | 2194-2339 | 6-28 | GACATCAGTAGATTGTACAATGT | AUUGUACAAUCUACUGAUGUC CAUCAGUAGAUUGUACAAUGU | **UA** | 20.4 °C | 20.3 °C |
| 85 | g85 | 2194-2339 | 9-31 | ATCAGTAGATTGTACAATGTACA | UACAUUGUACAAUCUACUGAU CAGUAGAUUGUACAAUGUACA | **UA** | 19.3 °C | 18.9 °C |
| 86 | g86 | 2194-2339 | 11-33 | CAGTAGATTGTACAATGTACATT | UGUACAUUGUACAAUCUACUG GUAGAUUGUACAAUGUACAUU | **U** | 14.6 °C | 13.4 °C |
| 87 | g87 | 2194-2339 | 14-36 | TAGATTGTACAATGTACATTTGT | AAAUGUACAUUGUACAAUCUA GAUUGUACAAUGUACAUUUGU | **UA** | 14.6 °C | 14.6 °C |
| 88 | g88 | 2194-2339 | 28-50 | TACATTTGTGGTGATTCAACTGA | AGUUGAAUCACCACAAAUGUA CAUUUGUGGUGAUUCAACUGA | **U** | 14.8 °C | 12.1 °C |
| 89 | g89 | 2194-2339 | 44-66 | CAACTGAATGCAGCAATCTTTTG | AAAGAUUGCUGCAUUCAGUUG ACUGAAUGCAGCAAUCUUUUG | **A** | 12.0 °C | 18.1 °C |
| 90 | g90 | 2194-2339 | 54-76 | CAGCAATCTTTTGTTGCAATATG | UAUUGCAACAAAAGAUUGCUG GCAAUCUUUUGUUGCAAUAUG | **URA** | 20.0 °C | 12.0 °C |
| 91 | g91 | 2194-2339 | 55-77 | AGCAATCTTTTGTTGCAATATGG | AUAUUGCAACAAAAGAUUGCU CAAUCUUUUGUUGCAAUAUGG | **UA** | 21.1 °C | 5.3 °C |
| 92 | g92 | 2194-2339 | 67-89 | TTGCAATATGGCAGTTTTTGTAC | ACAAAAACUGCCAUAUUGCAA GCAAUAUGGCAGUUUUUGUAC | **URA** | 5.6 °C | 5.6 °C |
| 93 | g93 | 2194-2339 | 68-90 | TGCAATATGGCAGTTTTTGTACA | UACAAAAACUGCCAUAUUGCA CAAUAUGGCAGUUUUUGUACA | **UR** | 5.6 °C | 12.6 °C |
| 94 | g94 | 2194-2339 | 75-97 | TGGCAGTTTTTGTACACAATTAA | AAUUGUGUACAAAAACUGCCA GCAGUUUUUGUACACAAUUAA | **UA** | 19.3 °C | 10.3 °C |
| 95 | g95 | 2194-2339 | 76-98 | GGCAGTTTTTGTACACAATTAAA | UAAUUGUGUACAAAAACUGCC CAGUUUUUGUACACAAUUAAA | **URA** | 12.1 °C | 3.2 °C |
| 96 | g96 | 2194-2339 | 77-99 | GCAGTTTTTGTACACAATTAAAC | UUAAUUGUGUACAAAAACUGC AGUUUUUGUACACAAUUAAAC | **R** | 6.9 °C | 5.6 °C |
| 97 | g97 | 2194-2339 | 78-100 | CAGTTTTTGTACACAATTAAACC | UUUAAUUGUGUACAAAAACUG GUUUUUGUACACAAUUAAACC | **URA** | -1.4 °C | 5.6 °C |
| 98 | g98 | 2194-2339 | 89-111 | CACAATTAAACCGTGCTTTAACT | UUAAAGCACGGUUUAAUUGUG CAAUUAAACCGUGCUUUAACU | **URA** | 19.7 °C | -9.7 °C |
| 99 | g99 | 2194-2339 | 109-131 | ACTGGAATAGCTGTTGAACAAGA | UUGUUCAACAGCUAUUCCAGU UGGAAUAGCUGUUGAACAAGA | **R** | 20.5 °C | 19.9 °C |
| 100 | g100 | 2194-2339 | 111-133 | TGGAATAGCTGTTGAACAAGACA | UCUUGUUCAACAGCUAUUCCA GAAUAGCUGUUGAACAAGACA | **UR** | 19.2 °C | 18.3 °C |
| 101 | g101 | 2194-2339 | 117-139 | AGCTGTTGAACAAGACAAAAACA | UUUUUGUCUUGUUCAACAGCU CUGUUGAACAAGACAAAAACA | **URA** | 14.9 °C | 20.5 °C |
| 102 | g102 | 2194-2339 | 119-141 | CTGTTGAACAAGACAAAAACACC | UGUUUUUGUCUUGUUCAACAG GUUGAACAAGACAAAAACACC | **UA** | 5.6 °C | 20.5 °C |
| 103 | g103 | 2534-2567 | 10-32 | ACCTCATTTGTGCACAAAAGTTT | ACUUUUGUGCACAAAUGAGGU CUCAUUUGUGCACAAAAGUUU | **UA** | 10.3 °C | 13.8 °C |
| 104 | g104 | 2534-2567 | 12-34 | CTCATTTGTGCACAAAAGTTTAA | AAACUUUUGUGCACAAAUGAG CAUUUGUGCACAAAAGUUUAA | **UA** | 3.2 °C | 12.1 °C |
| 105 | g105 | 2676-2720 | 19-41 | TGCTATGCAAATGGCTTATAGGT | CUAUAAGCCAUUUGCAUAGCA CUAUGCAAAUGGCUUAUAGGU | **R** | 14.9 °C | 21.1 °C |
| 106 | g106 | 2676-2720 | 23-45 | ATGCAAATGGCTTATAGGTTTAA | AAACCUAUAAGCCAUUUGCAU GCAAAUGGCUUAUAGGUUUAA | **URA** | 18.5 °C | 17.7 °C |
| 107 | g107 | 2722-2785 | 6-28 | TGGAGTTACACAGAATGTTCTCT | AGAACAUUCUGUGUAACUCCA GAGUUACACAGAAUGUUCUCU | **UA** | 14.8 °C | 19.0 °C |
| 108 | g108 | 2722-2785 | 12-34 | TACACAGAATGTTCTCTATGAGA | UCAUAGAGAACAUUCUGUGUA CACAGAAUGUUCUCUAUGAGA | **URA** | 17.8 °C | 19.2 °C |
| 109 | g109 | 2722-2785 | 14-36 | CACAGAATGTTCTCTATGAGAAC | UCUCAUAGAGAACAUUCUGUG CAGAAUGUUCUCUAUGAGAAC | **UA** | 17.8 °C | 19.2 °C |
| 110 | g110 | 2722-2785 | 15-37 | ACAGAATGTTCTCTATGAGAACC | UUCUCAUAGAGAACAUUCUGU AGAAUGUUCUCUAUGAGAACC | **R** | 21.4 °C | 14.8 °C |
| 111 | g111 | 2722-2785 | 22-44 | GTTCTCTATGAGAACCAAAAATT | UUUUUGGUUCUCAUAGAGAAC UCUCUAUGAGAACCAAAAAUU | **R** | 18.8 °C | 17.8 °C |
| 112 | g112 | 2722-2785 | 23-45 | TTCTCTATGAGAACCAAAAATTG | AUUUUUGGUUCUCAUAGAGAA CUCUAUGAGAACCAAAAAUUG | **UA** | 11.5 °C | 21.4 °C |
| 113 | g113 | 2829-2903 | 29-51 | AACCAAAATGCACAAGCTTTAAA | UAAAGCUUGUGCAUUUUGGUU CCAAAAUGCACAAGCUUUAAA | **URA** | 17.0 °C | 4.2 °C |
| 114 | g114 | 2829-2903 | 30-52 | ACCAAAATGCACAAGCTTTAAAC | UUAAAGCUUGUGCAUUUUGGU CAAAAUGCACAAGCUUUAAAC | **URA** | 18.3 °C | 14.0 °C |
| 115 | g115 | 2829-2903 | 31-53 | CCAAAATGCACAAGCTTTAAACA | UUUAAAGCUUGUGCAUUUUGG AAAAUGCACAAGCUUUAAACA | **R** | 13.7 °C | 20.0 °C |
| 116 | g116 | 2829-2903 | 43-65 | AGCTTTAAACACGCTTGTTAAAC | UUAACAAGCGUGUUUAAAGCU CUUUAAACACGCUUGUUAAAC | **UA** | 11.8 °C | 0.0 °C |
| 117 | g117 | 2829-2903 | 44-66 | GCTTTAAACACGCTTGTTAAACA | UUUAACAAGCGUGUUUAAAGC UUUAAACACGCUUGUUAAACA | **R** | 7.2 °C | 7.2 °C |
| 118 | g118 | 2829-2903 | 46-68 | TTTAAACACGCTTGTTAAACAAC | UGUUUAACAAGCGUGUUUAAA UAAACACGCUUGUUAAACAAC | **R** | 8.2 °C | 19.8 °C |
| 119 | g119 | 2905-2939 | 6-28 | TGGTGCAATTTCAAGTGTTTTAA | AAAACACUUGAAAUUGCACCA GUGCAAUUUCAAGUGUUUUAA | **UA** | 17.8 °C | 20.0 °C |
| 120 | g120 | 2905-2939 | 7-29 | GGTGCAATTTCAAGTGTTTTAAA | UAAAACACUUGAAAUUGCACC UGCAAUUUCAAGUGUUUUAAA | **R** | 13.3 °C | 14.0 °C |
| 121 | g121 | 2905-2939 | 8-30 | GTGCAATTTCAAGTGTTTTAAAT | UUAAAACACUUGAAAUUGCAC GCAAUUUCAAGUGUUUUAAAU | **URA** | 7.2 °C | 7.4 °C |
| 122 | g122 | 2905-2939 | 9-31 | TGCAATTTCAAGTGTTTTAAATG | UUUAAAACACUUGAAAUUGCA CAAUUUCAAGUGUUUUAAAUG | **URA** | 0.0 °C | 7.4 °C |
| 123 | g123 | 2905-2939 | 10-32 | GCAATTTCAAGTGTTTTAAATGA | AUUUAAAACACUUGAAAUUGC AAUUUCAAGUGUUUUAAAUGA | **R** | -9.1 °C | 7.4 °C |
| 124 | g124 | 2905-2939 | 12-34 | AATTTCAAGTGTTTTAAATGATA | UCAUUUAAAACACUUGAAAUU UUUCAAGUGUUUUAAAUGAUA | **R** | -1.4 °C | 19.2 °C |
| 125 | g125 | 2992-3110 | 24-46 | GACATATGTGACTCAACAATTAA | AAUUGUUGAGUCACAUAUGUC CAUAUGUGACUCAACAAUUAA | **UA** | 12.1 °C | 13.3 °C |
| 126 | g126 | 2992-3110 | 25-47 | ACATATGTGACTCAACAATTAAT | UAAUUGUUGAGUCACAUAUGU AUAUGUGACUCAACAAUUAAU | **R** | 5.3 °C | 21.5 °C |
| 127 | g127 | 2992-3110 | 73-95 | GCTAATCTTGCTGCTACTAAAAT | UUUAGUAGCAGCAAGAUUAGC UAAUCUUGCUGCUACUAAAAU | **R** | 11.3 °C | 12.0 °C |
| 128 | g128 | 3385-3427 | 12-34 | TGTCAACAACACAGTTTATGATC | UCAUAAACUGUGUUGUUGACA UCAACAACACAGUUUAUGAUC | **R** | 6.9 °C | 19.3 °C |
| 129 | g129 | 3385-3427 | 13-35 | GTCAACAACACAGTTTATGATCC | AUCAUAAACUGUGUUGUUGAC CAACAACACAGUUUAUGAUCC | **UA** | 8.9 °C | 19.3 °C |
| 130 | g130 | 3602-3647 | 8-30 | TGGAAAGTATGAGCAGTATATAA | AUAUACUGCUCAUACUUUCCA GAAAGUAUGAGCAGUAUAUAA | **URA** | 13.1 °C | 4.6 °C |
| 131 | g131 | 3602-3647 | 9-31 | GGAAAGTATGAGCAGTATATAAA | UAUAUACUGCUCAUACUUUCC AAAGUAUGAGCAGUAUAUAAA | **R** | 6.1 °C | 11.6 °C |
| 132 | g132 | 3602-3647 | 10-32 | GAAAGTATGAGCAGTATATAAAA | UUAUAUACUGCUCAUACUUUC AAGUAUGAGCAGUAUAUAAAA | **R** | 2.8 °C | 20.3 °C |
| 133 | g133 | 3602-3647 | 11-33 | AAAGTATGAGCAGTATATAAAAT | UUUAUAUACUGCUCAUACUUU AGUAUGAGCAGUAUAUAAAAU | **R** | -5.9 °C | 20.3 °C |
| 134 | g134 | 3649-3713 | 18-40 | TGGCTTGATTGCCATAGTAATGG | AUUACUAUGGCAAUCAAGCCA GCUUGAUUGCCAUAGUAAUGG | **UA** | 6.3 °C | 12.0 °C |
| 135 | g135 | 3649-3713 | 28-50 | GCCATAGTAATGGTGACAATTAT | AAUUGUCACCAUUACUAUGGC CAUAGUAAUGGUGACAAUUAU | **UA** | 20.5 °C | 6.3 °C |
| 136 | g136 | 3649-3713 | 29-51 | CCATAGTAATGGTGACAATTATG | UAAUUGUCACCAUUACUAUGG AUAGUAAUGGUGACAAUUAUG | **R** | 14.8 °C | 6.3 °C |
| 137 | g137 | 3649-3713 | 30-52 | CATAGTAATGGTGACAATTATGC | AUAAUUGUCACCAUUACUAUG UAGUAAUGGUGACAAUUAUGC | **R** | 6.9 °C | 11.6 °C |
