## Supplementary Table 10 for "Computational study and design of effective siRNAs to silence structural proteins associated genes of Indian SARS-CoV-2 strains"

**Supplementary Table 10.** Effective siRNAs against envelope gene with GC%, free energy of folding and free energy of binding with target.

| **S.no** | **Alias** | **GC%** | **Free energy of folding** | **Free energy of binding** | **Tm (Conc)** | **Tm (Cp)** |
| --- | --- | --- | --- | --- | --- | --- |
| 1 | e1 | 31% | 1.7 | -29.1 | 78.4 | 79.6 |
| 2 | e2 | 29% | 1.7 | -29 | 82.6 | 83.9 |
| 3 | e3 | 29% | 1.7 | -28.6 | 82.6 | 84.3 |
| 4 | e4 | 29% | 1.8 | -27.9 | 82 | 83.6 |
| 5 | e5 | 38% | 1.7 | -34.1 | 85 | 86 |
| 6 | e6 | 21% | 1.3 | -25.8 | 76.4 | 77.6 |
| 7 | e7 | 26% | 1.4 | -28 | 80.4 | 81.3 |
