## Supplementary Table 11 for "Computational study and design of effective siRNAs to silence structural proteins associated genes of Indian SARS-CoV-2 strains"

**Supplementary Table 11.** Effective siRNAs against membrane gene with GC%, free energy of folding and free energy of binding with target.

| **S.no** | **Alias** | **GC%** | **Free energy of folding** | **Free energy of binding** | **Tm (Conc)** | **Tm (Cp)** |
| --- | --- | --- | --- | --- | --- | --- |
| 1 | m1 | 38% | 1.8 | -31.3 | 87.5 | 88.7 |
| 2 | m2 | 33% | 1.6 | -33.5 | 86.3 | 86.9 |
| 3 | m3 | 33% | 2 | -30.8 | 87.4 | 88.6 |
| 4 | m4 | 21% | 1.8 | -28.4 | 74.9 | 76.2 |
| 5 | m5 | 19% | 1.8 | -27.5 | 74.4 | 76.1 |
| 6 | m6 | 17% | 1.8 | -24.8 | 73.9 | 75.7 |
| 7 | m7 | 17% | 1.8 | -23.9 | 71.6 | 73.2 |
| 8 | m8 | 17% | 1.8 | -23.7 | 70.2 | 71.8 |
| 9 | m9 | 10% | 1.8 | -20.9 | 68.5 | 70.2 |
| 10 | m10 | 10% | 1.7 | -20.9 | 66.8 | 68.4 |
