## Supplementary Table 12 for "Computational study and design of effective siRNAs to silence structural proteins associated genes of Indian SARS-CoV-2 strains"

**Supplementary Table 12.** Effective siRNAs against nucleocapsid phosphoprotein gene with GC%, free energy of folding and free energy of binding with target.

| **S.no** | **Alias** | **GC%** | **Free energy of folding** | **Free energy of binding** | **Tm (Conc)** | **Tm (Cp)** |
| --- | --- | --- | --- | --- | --- | --- |
| 1 | n1 | 40% | 1.8 | -35.8 | 86.2 | 87.6 |
| 2 | n2 | 38% | 1.7 | -34.9 | 88.9 | 90.3 |
| 3 | n3 | 36% | 1.9 | -32.5 | 87.9 | 89.4 |
