## Supplementary Table 13 for "Computational study and design of effective siRNAs to silence structural proteins associated genes of Indian SARS-CoV-2 strains"

**Supplementary Table 13.** Effective siRNAs against surface glycoprotein gene with GC%, free energy of folding and free energy of binding with target.

| **S.no** | **Alias** | **GC%** | **Free energy of folding** | **Free energy of binding** | **Tm (Conc)** | **Tm (Cp)** |
| --- | --- | --- | --- | --- | --- | --- |
| 1 | g1 | 29% | 1.8 | -28.9 | 81.1°C | 81.9°C |
| 2 | g2 | 26% | 1.7 | -29.7 | 78.8°C | 80.1°C |
| 3 | g3 | 26% | 1.6 | -28.9 | 75.6°C | 77.0°C |
| 4 | g4 | 21% | 1.6 | -27.4 | 73.7°C | 75.1°C |
| 5 | g5 | 21% | 1.6 | -25 | 72.6°C | 73.9°C |
| 6 | g6 | 31% | 1.8 | -31.3 | 78.2°C | 79.4°C |
| 7 | g7 | 29% | 1.7 | -30.4 | 78.2°C | 79.3°C |
| 8 | g8 | 24% | 1.7 | -27 | 77.6°C | 79.2°C |
| 9 | g9 | 21% | 1.7 | -25.3 | 77.8°C | 79.3°C |
| 10 | g10 | 29% | 1.8 | -29.7 | 81.2°C | 82.6°C |
| 11 | g11 | 26% | 1.8 | -29.6 | 77.2°C | 78.7°C |
| 12 | g12 | 24% | 1.8 | -29.2 | 74.0°C | 75.3°C |
| 13 | g13 | 21% | 1.8 | -26.5 | 73.2°C | 74.9°C |
| 14 | g14 | 19% | 1.8 | -25.4 | 76.9°C | 78.5°C |
| 15 | g15 | 19% | 1.8 | -25.3 | 75.4°C | 76.9°C |
| 16 | g16 | 19% | 1.8 | -25.5 | 74.3°C | 75.4°C |
| 17 | g17 | 17% | 1.8 | -26 | 73.3°C | 74.7°C |
| 18 | g18 | 17% | 1.8 | -25.3 | 74.1°C | 75.6°C |
| 19 | g19 | 14% | 1.8 | -23.3 | 74.5°C | 76.0°C |
| 20 | g20 | 14% | 1.8 | -23.3 | 70.5°C | 71.8°C |
| 21 | g21 | 12% | 1.8 | -23.5 | 67.9°C | 69.5°C |
| 22 | g22 | 14% | 1.7 | -22.5 | 67.3°C | 69.0°C |
| 23 | g23 | 12% | 1.7 | -20.5 | 65.6°C | 67.2°C |
| 24 | g24 | 10% | 1.6 | -21.6 | 66.2°C | 67.7°C |
| 25 | g25 | 12% | 1.6 | -20.6 | 63.4°C | 65.2°C |
| 26 | g26 | 17% | 1.8 | -23.5 | 71.2°C | 72.8°C |
| 27 | g27 | 31% | 1.5 | -29.9 | 82.7°C | 83.4°C |
| 28 | g28 | 24% | 1.7 | -28.3 | 80.3°C | 81.2°C |
| 29 | g29 | 21% | 1.8 | -27.7 | 79.3°C | 80.4°C |
| 30 | g30 | 19% | 1.8 | -25.7 | 79.3°C | 80.3°C |
| 31 | g31 | 21% | 1.8 | -25.9 | 77.7°C | 78.4°C |
| 32 | g32 | 21% | 1.8 | -25.9 | 76.3°C | 76.9°C |
| 33 | g33 | 21% | 1.8 | -28 | 76.3°C | 77.3°C |
| 34 | g34 | 21% | 1.5 | -25.9 | 75.9°C | 77.1°C |
| 35 | g35 | 21% | 1.6 | -27.8 | 76.3°C | 76.9°C |
| 36 | g36 | 38% | 1.9 | -36.1 | 88.9°C | 89.3° |
| 37 | g37 | 36% | 1.9 | -33.6 | 88.2°C | 89.0°C |
| 38 | g38 | 33% | 1.8 | -31.2 | 87.9°C | 89.1°C |
| 39 | g39 | 33% | 1.8 | -31.7 | 83.3°C | 84.7°C |
| 40 | g40 | 29% | 2 | -29.9 | 79.7°C | 80.9°C |
| 41 | g41 | 26% | 1.9 | -27.8 | 79.2°C | 80.5°C |
| 42 | g42 | 26% | 1.6 | -26.8 | 80.2°C | 81.6°C |
| 43 | g43 | 31% | 1.3 | -30.6 | 80.3°C | 81.6°C |
| 44 | g44 | 29% | 1.4 | -30.3 | 78.7°C | 80.2°C |
| 45 | g45 | 26% | 1.7 | -27.9 | 78.5°C | 79.9°C |
| 46 | g46 | 24% | 1.6 | -28.2 | 77.3°C | 78.5°C |
| 47 | g47 | 26% | 1.6 | -28.2 | 80.4°C | 79.1°C |
| 48 | g48 | 29% | 1.8 | -32.8 | 81.4°C | 82.5°C |
| 49 | g49 | 26% | 1.8 | -30.8 | 80.8°C | 81.5°C |
| 50 | g50 | 21% | 1.8 | -28.9 | 81.0°C | 82.4°C |
| 51 | g51 | 24% | 1.7 | -27.9 | 81.0°C | 82.3°C |
| 52 | g52 | 31% | 1.8 | -32.2 | 81.9°C | 83.3°C |
| 53 | g53 | 26% | 1.8 | -30.3 | 78.9°C | 80.2°C |
| 54 | g54 | 24% | 1.8 | -29.3 | 77.6°C | 79.1°C |
| 55 | g55 | 14% | 1.8 | -23.2 | 71.0°C | 72.5°C |
| 56 | g56 | 14% | 1.8 | -23.8 | 65.4°C | 67.0°C |
| 57 | g57 | 14% | 1.7 | -23.3 | 63.1°C | 64.7°C |
| 58 | g58 | 14% | 1.7 | -23.6 | 68.7°C | 70.3°C |
| 59 | g59 | 21% | 1.7 | -25.9 | 78.4°C | 79.8°C |
| 60 | g60 | 21% | 1.7 | -25.7 | 78.4°C | 79.9°C |
| 61 | g61 | 33% | 1.6 | -31.7 | 68.3°C | 69.0°C |
| 62 | g62 | 29% | 1.6 | -30 | 79.9°C | 81.0°C |
| 63 | g63 | 29% | 1.6 | -29.7 | 77.3°C | 78.6°C |
| 64 | g64 | 31% | 1.6 | -29.6 | 78.8°C | 80.3°C |
| 65 | g65 | 33% | 1.5 | -31.3 | 83.4°C | 83.5°C |
| 66 | g66 | 29% | 1.6 | -30.3 | 80.2°C | 81.2°C |
| 67 | g67 | 29% | 1.6 | -29.3 | 80.1°C | 81.5°C |
| 68 | g68 | 38% | 1.8 | -33.3 | 91.1°C | 92.1°C |
| 69 | g69 | 36% | 1.8 | -31.6 | 91.1°C | 92.1°C |
| 70 | g70 | 31% | 1.9 | -32.4 | 83.6°C | 84.6°C |
| 71 | g71 | 29% | 1.9 | -31.4 | 81.3°C | 82.6°C |
| 72 | g72 | 29% | 1.9 | -29 | 79.2°C | 80.5°C |
| 73 | g73 | 26% | 1.6 | -27.6 | 76.1°C | 77.2°C |
| 74 | g74 | 26% | 1.8 | -26.7 | 78.2°C | 79.5°C |
| 75 | g75 | 32% | 1.7 | -28.8 | 79.2°C | 80.7°C |
| 76 | g76 | 33% | 1.6 | -29.6 | 80.2°C | 81.5°C |
| 77 | g77 | 33% | 1.9 | -31.6 | 84.5°C | 85.6°C |
| 78 | g78 | 33% | 1.9 | -31.6 | 83.3°C | 84.5°C |
| 79 | g79 | 36% | 2 | -31.5 | 82.9°C | 84.2°C |
| 80 | g80 | 43% | 1.9 | -34.7 | 89.7°C | 91.0°C |
| 81 | g81 | 36% | 1.9 | -32.7 | 83.7°C | 84.1°C |
| 82 | g82 | 40% | 1.8 | -35.7 | 87.4°C | 88.2°C |
| 83 | g83 | 38% | 1.5 | -34.9 | 83.5°C | 84.7°C |
| 84 | g84 | 33% | 1.7 | -32.4 | 82.6°C | 84.0°C |
| 85 | g85 | 31% | 1.6 | -30.6 | 81.1°C | 82.2°C |
| 86 | g86 | 31% | 2 | -32 | 81.1°C | 82.0°C |
| 87 | g87 | 26% | 2 | -28.1 | 78.6°C | 79.4°C |
| 88 | g88 | 36% | 1.7 | -32.2 | 84.5°C | 85.8°C |
| 89 | g89 | 38% | 1.9 | -32.8 | 86.5°C | 87.5°C |
| 90 | g90 | 33% | 2 | -31 | 80.9°C | 81.7°C |
| 91 | g91 | 31% | 2 | -29.6 | 77.7°C | 78.8°C |
| 92 | g92 | 36% | 2 | -31.6 | 83.4°C | 83.8°C |
| 93 | g93 | 33% | 1.9 | -31.8 | 81.9°C | 82.6°C |
| 94 | g94 | 31% | 1.5 | -31.5 | 81.2°C | 82.5°C |
| 95 | g95 | 29% | 1.7 | -31.3 | 77.5°C | 78.8°C |
| 96 | g96 | 26% | 1.7 | -28.9 | 76.2°C | 77.4°C |
| 97 | g97 | 26% | 1.7 | -26.1 | 75.6°C | 76.8°C |
| 98 | g98 | 33% | 1.9 | -30 | 83.5°C | 84.7°C |
| 99 | g99 | 38% | 1.6 | -34.5 | 87.3°C | 88.7°C |
| 100 | g100 | 38% | 1.6 | -34.4 | 85.1°C | 86.3°C |
| 101 | g101 | 33% | 1.5 | -31.2 | 83.8°C | 85.1°C |
| 102 | g102 | 36% | 1.5 | -30.5 | 81.0°C | 82.0°C |
| 103 | g103 | 36% | 1.1 | -33.2 | 84.9°C | 84.9°C |
| 104 | g104 | 31% | 1.5 | -30.6 | 82.3°C | 82.6°C |
| 105 | g105 | 38% | 1.5 | -34.8 | 85.1°C | 86.0°C |
| 106 | g106 | 33% | 1.9 | -32.1 | 85.4°C | 86.6°C |
| 107 | g107 | 38% | 1.6 | -34.1 | 83.4°C | 84.2°C |
| 108 | g108 | 36% | 1.9 | -33.2 | 84.2°C | 83.7°C |
| 109 | g109 | 38% | 1.9 | -34.7 | 84.4°C | 84.0°C |
| 110 | g110 | 36% | 1.4 | -32.7 | 84.2°C | 84.0°C |
| 111 | g111 | 31% | 2.9 | -31.6 | 86.9°C | 87.5°C |
| 112 | g112 | 31% | 1.9 | -29.5 | 85.5°C | 86.0°C |
| 113 | g113 | 33% | 1.9 | -31.5 | 84.7°C | 85.7°C |
| 114 | g114 | 33% | 1.8 | -31.5 | 83.5°C | 84.6°C |
| 115 | g115 | 31% | 1.8 | -30.4 | 83.4°C | 85.1°C |
| 116 | g116 | 33% | 1.7 | -30.9 | 84.0°C | 84.7°C |
| 117 | g117 | 31% | 1.5 | -29.9 | 83.9°C | 84.9°C |
| 118 | g118 | 31% | 1.3 | -28.5 | 83.5°C | 84.2°C |
| 119 | g119 | 31% | 1.8 | -31.3 | 80.6°C | 82.2°C |
| 120 | g120 | 29% | 1.8 | -31.1 | 79.1°C | 80.7°C |
| 121 | g121 | 26% | 1.8 | -28.7 | 77.7°C | 79.1°C |
| 122 | g122 | 24% | 1.6 | -26.7 | 74.2°C | 75.6°C |
| 123 | g123 | 21% | 1.6 | -26.3 | 73.4°C | 75.1°C |
| 124 | g124 | 19% | 1.5 | -24.7 | 76.1°C | 77.5°C |
| 125 | g125 | 31% | 1.6 | -32 | 82.8°C | 84.1°C |
| 126 | g126 | 26% | 1.7 | -30.5 | 82.3°C | 83.8°C |
| 127 | g127 | 33% | 1.7 | -34.3 | 88.5°C | 89.8°C |
| 128 | g128 | 33% | 1.6 | -32 | 82.1°C | 83.3°C |
| 129 | g129 | 36% | 1.8 | -31 | 80.6°C | 82.0°C |
| 130 | g130 | 31% | 1.8 | -33.1 | 83.6°C | 84.9°C |
| 131 | g131 | 29% | 1.8 | -32.9 | 82.9°C | 84.3°C |
| 132 | g132 | 26% | 1.8 | -30.5 | 83.2°C | 84.3°C |
| 133 | g133 | 24% | 1.8 | -28.5 | 82.4°C | 83.4°C |
| 134 | g134 | 40% | 2 | -34.8 | 86.1°C | 87.1°C |
| 135 | g135 | 33% | 2 | -34.2 | 83.5°C | 84.6°C |
| 136 | g136 | 31% | 2 | -32 | 82.7°C | 84.0°C |
| 137 | g137 | 31% | 1.7 | -29.9 | 82.2°C | 83.5°C |
