## Supplementary Table 14 for "Computational study and design of effective siRNAs to silence structural proteins associated genes of Indian SARS-CoV-2 strains"

**Supplementary Table 14.** siRNA and Ago2 interacting residues and bond category.

| **S.no** | **siRNA in Ago2-siRNA complex** | **Guide strand (5' →3')** | **HDock Docking score** | **Interaction Category** | | | | **Interacting Residues** | |
| --- | --- | --- | --- | --- | --- | --- | --- | --- | --- |
|  |  |  |  | **Salt bridge (Hydrogen Bond+Electrostatic)** | **Hydrogen bonds** | **Electrostatic bonds** | **Hydrophobic bonds** | **Amino acid** | **Total count** |
| 1 | e5 | UGUAACUAGCAAGAAUACCAC | -322.34 | 7 | 16 | 8 | 5 | TYR279, THR222, SER220, PRO67, PRO63, LYS65, LYS62, LYS278, LYS129, LYS112, GLY674, GLU673, ARG97, ARG710, ARG68, ARG635, ARG351, ARG280, ARG179, ARG167, GLN160, VAL177, GLU58 | 23 |
| 2 | m1 | UUUUUAAGCUCUUCAACGGUA | -324.56 | 3 | 13 | 11 | 6 | ARG179, ARG351, ARG366, ARG635, ARG710, ARG711, ARG761, ARG792, ARG795, ARG812, ASN359, ASN99, GLY674, HIS807, LYS550, LYS709, SER362, SER672, THR556, THR559, LEU808, PHE811, PHE858, SER220, HIS753, ILE365, SER36 | 27 |
| 3 | m2 | UGUUCAAGGAGCUUUUUAAGC | -348.34 | 3 | 15 | 8 | 3 | ARG255, ARG837, ARG97, LYS263, ARG286, LYS607, ARG635, LYS98, ASN99, ARG280, ASN283, GLN633, ARG710, GLU333, HIS336, SER672, ASP95, GLU637, LYS65, GLN675 | 20 |
| 4 | n1 | UUUCUUAGUGACAGUUUGGCC | -335.64 | 1 | 15 | 7 | 0 | ARG710, LYS98, ARG351, ARG635, ASN99, GLN160, GLN708, GLU637, GLY674, ASP95, HIS681, THR599, HIS600, GLY670, GLN675, ARG97 | 16 |
| 5 | n2 | AUUUCCUUGGGUUUGUUCUGG | -294.72 | 1 | 10 | 9 | 5 | ARG277, ARG69, CYS272, GLN332, HIS316, HIS336, LYS129, LYS335, LYS65, THR337, TYR279, TYR311, PRO63, LYS62 | 14 |
| 6 | g36 | UUUCUACAGUACCAAAAGGGC | -354.19 | 1 | 12 | 7 | 2 | ARG179, ARG351, ARG688, ARG710, ARG761, GLN633, GLN636, GLN640, GLN678, HIS681, HIS711, HIS712, LYS696, SER672, ILE638, THR222 | 16 |
| 7 | g82 | AUAACUAGCGCAUAUACCUGC | -290.15 | 1 | 21 | 10 | 3 | ARG97, LYS65, ARG68, LYS98, ARG179, ARG351, ARG635, ARG710, GLN633, HIS634, GLN675, GLN678, GLU64, GLY674, PRO602, GLN636, SER672, LYS607, ILE353 | 19 |
| 8 | g83 | ACAAUCUACUGAUGUCUUGGU | -326.1 | 1 | 12 | 4 | 5 | ARG647, ARG658, ARG854, GLY514, HIS849, LYS655, SER656, GLN850, LYS844, LEU515, ILE651, GLN516 | 12 |
| 9 | g105 | CUAUAAGCCAUUUGCAUAGCA | -312.45 | 2 | 14 | 3 | 3 | ARG280, ARG351, ARG68, LYS98, GLN633, ARG635, ARG710, HIS839, GLN678, GLN332, GLU333, SER672, ASP641, HIS681, HIS682, LYS65, ALA644 | 17 |
| 10 | g134 | AUUACUAUGGCAAUCAAGCCA | -338.34 | 0 | 14 | 3 | 2 | LYS98, ARG167, ARG68, ARG97, ASN99, GLN160, HIS681, GLN677, ASP163, GLN636, LYS693, PHE156 | 12 |
| 11 | g80 | AAUAAACACGCCAAGUAGGAG | -358.89 | 4 | 8 | 10 | 2 | ARG255, ARG280, LYS354, ARG761, ARG286, ARG351, ARG710, ARG714, ARG792, TYR790, GLY331, GLU333, LYS263, TYR804, HIS712, ILE756, ALA369 | 17 |
| 12 | g109 | UCUCAUAGAGAACAUUCUGUG | -319.69 | 3 | 12 | 7 | 2 | ARG68, ARG97, LYS65, LYS607, HIS634, GLN678, GLU637, GLU64, GLN64, ARG635, GLN636, HIS682 | 12 |
